# Novel Synthetic Polymyxin Variants Inspired by Newly Uncovered Natural Sequences Explored as Potential Antibiotics

**DOI:** 10.64898/2026.09.25.754415

**Authors:** Walliyulahi Ajibola, Roland Tengölics, Bakhtiyar Mahmood, Annamária Marton, Éva Hunyadi-Gulyás, Csaba Vizler, József Sóki, Zsuzsanna Darula, Tamás Fehér

## Abstract

The incidence of infections caused by multidrug-resistant bacterial agents has increased at an alarming rate worldwide. With the aim of identifying novel antimicrobial peptides (AMPs), we screened bacteria isolated from various environmental samples. Our hypothesis was that the sequence space of natural AMPs belonging to known AMP classes is far from saturation. We used a classical pipeline of bacterial culturing, overlay assays, extraction and fractionation to purify AMPs for identification. Using LC/MS analysis, the structure of AMPs isolated from a strain of *Paenibacillus* was narrowed down and a new subclass of polymyxins was uncovered. These harbor three aliphatic residues within the cyclic C-terminal, and in certain cases Ser replaces Thr at position A2. Four discrete polymyxins fitting the new class, but not identical to the natural polymyxins were synthesized and tested against 29 human pathogenic bacteria. Each displayed an antibacterial spectrum different from that of colistin, with the best candidate surpassing it in potency against ten bacterial strains, but underperforming against three others. These results demonstrate that current screening of natural bacterial isolates can still permit the design of novel antimicrobial peptide variants without the substantial threat of diminishing returns.

## Introduction

Bacterial infections in humans caused by antimicrobial resistant (AMR) pathogens pose an increasing threat worldwide. In 2021, a total of 4.7 million deaths were associated with AMR, and 1.14 million deaths were directly attributed to infections caused by AMR bacteria (Naghavi et al., 2024). From 1990 to 2021, the number of deaths attributable to methicillin-resistant *Staphylococcus aureus* (MRSA) and carbapenem-resistant Gram-negative bacteria increased by 128% and 70%, respectively (Naghavi et al., 2024). The total hospital cost associated with AMR was estimated to gross US$ 698 billion in 2019 (Naylor et al., 2025). Alarmingly, the global prevalence of AMR has increased in 40% of the pathogen– antibiotic combinations monitored between 2018 and 2023 (Tosas Auguet et al., 2025).

Several strategies have been applied to address the unfavorable trends described above. These include the foundation of a global surveillance system on antibiotic resistance, initiation of national action plans on antibiotic use in health care and agriculture, improving education and public awareness, and last but not least, developing novel antimicrobial therapies (Karnwal et al., 2025). Unfortunately, there has been a dramatic decline in the discovery of antibiotics of novel class, with just two antibiotics of new structural types (daptomycin and linezolid) reaching the market in the last 40 years (Cochrane & Vederas, 2016). This fact stands in stark contrast with the opinion that nature still provides a vast and untapped resource of antimicrobial compounds (Karnwal & Malik, 2024).

Despite the possibilities provided by the *de novo* design of completely synthetic antibiotics, the majority of today’s antimicrobial therapies are based on natural compounds, or on modifications thereof (Demain & Fang, 2000). The growing repertoire of compounds originating from microbiota (bacteria (Yi et al., 2026), fungi (Conrado et al., 2022) and protists (Senra, 2023)), plants (e.g. algae (Matin et al., 2024), medicinal plants (Marok et al., 2024)) and animals (e.g. mussels (Roch et al., 2008), sponges (Liu et al., 2024), bees, scorpions, snakes (Rabea et al., 2025)) promises to provide potential therapeutics against human, plant and veterinary pathogens in the future. The majority of the antibiotics in use today, or undergoing discovery, originate from bacteria, namely the *Streptomyces* species (Donald et al., 2022). The common rediscovery of antimicrobials extracted from bacterial samples has prompted researchers to shift their focus to previously under-explored habitats. In the course of this initiative, microbiomes originating from plants (Romero-Severson et al., 2021), animals (Oyama et al., 2017) and humans (Zipperer et al., 2016) have contributed to the available repertoire of experimentally validated antimicrobials. Even within this versatile resource of microbiota however, soil remains the most important microbiome reservoir for the search of secondary metabolites (Timmis & Ramos, 2021).

The oldest, most straightforward and historically most often applied strategy to isolate novel antimicrobial compounds is via functional screening of natural samples. This can take place either using classical inhibitory zone detection experiments (i.e. agar diffusion tests), or a number of evolved techniques including the resazurin assay, TLC bioautography, flow cytometry, bioluminescence or impedance measurement (Hossain, 2024). The improvement on gene annotation techniques later allowed the direct analysis of the DNA sequences of candidate microbiota, often referred to as genome mining. The majority of the microbial world, however, has not been cultured under laboratory conditions, which brought the advent of metagenomics; this is the identification of genes responsible for secondary metabolite production in DNA samples without mandatory knowledge of the producer strain itself, relying either on functional tests (Coughlan et al., 2015) or on comparative studies (Weisberg et al., 2026). Recently, genome mining has been combined with metabolomics to link secondary metabolites to their biosynthetic gene clusters, thereby accelerating de-replication and hence the discovery of novel compounds (Amoutzias et al., 2016). Currently, yet another shift in paradigm can be witnessed as AI-supported genome mining methods aiming to discover novel bioactive products become increasingly widespread (Wan et al., 2024).

With the extended repository of techniques used to screen, discover and exploit naturally derived antimicrobials, one might perceive that natural sources of such compounds are overmined. This work was, in part, initiated to test whether classical functional screens of environmental bacteria can still yield rewarding discoveries, or diminishing returns now make this approach unprofitable.

We report here the functional screening of numerous environmentally derived culturable bacteria, supported by fractionation, LC/MS-MS analysis, genome sequencing and peptide synthesis. Applying these tools and isolating an antimicrobial-producing *Paenibacillus* strain, we found a gene cluster encoding the synthesis of full length polymyxins harboring either three Leu/Ile residues or one Val and two Leu/Ile residues within their heptameric ring. We also detected natural polymyxins harboring a Ser residue at position A2. This paper describes several synthetic polymyxins inspired by but not identical to the natural polymyxins detected in our biological sample.

## Methods and Materials

### Strains and plasmids

*Escherichia coli* strains MDS42 (Pósfai et al., 2006) and DH10B ((Durfee et al., 2008)) were used for routine cloning purposes. *E. coli* Nissle 1917 (Cress et al., 2013), *E. coli* BAP1 (Pfeifer et al., 2001) and *Bacillus subtilis* 168 (Kunst et al., 1997) were used for heterologous expression of biosynthetic gene clusters (BGCs). *E. coli* strain BAP1 was obtained from Kerafast Inc. (Boston, MA, USA). The *Saccharomyces cerevisiae* YMH157 strain (a kind gift of Professor Harmit Malik, Fred Hutchinson Cancer Research Center, Seattle, USA) was used for yeast-based plasmid assembly and cloning.

Plasmid pJOE8999 (Altenbuchner, 2016) was a kind gift of Dr. Josef Altenbuchner. pJOE26_16 was made in two steps. First, the two homology arms were amplified from *Paenibacillus* ISO4 genomic DNA by PCR using primer pairs Xba26_16Up_F + 26_16Up_R and 26_16Dn_F + Xba26_16Dn_R (Supplementary Table S1), overlapped in overlap-extension PCR, digested by XbaI and cloned into the XbaI site of pJOE8999. Second, oligonucleotides sg26_16F and sg26_16R were hybridized and cloned into the BsaI site.

### Chemicals and media

Antibiotics were purchased from Sigma-Aldrich (St. Louis, MO, USA) and were used in the following concentrations kanamycin (Km): 25 µg/mL, chloramphenicol (Cm): 25 µg/mL, tetracycline (Tc): 10 µg/mL. Plasmid preparations were made using the Thermo Scientific GeneJET Plasmid Miniprep kit (Thermo Fisher Scientific Baltics UAB, Vilnius, Lithuania). For the preparation of bacterial artificial chromosomes (BACs), the Zymo Research ZR BAC DNA Miniprep kit (Zymo Research, Irvine, CA, USA) or the NucleoBond Extra Midi Plus Kit from Macherey Nagel (Düren, Germany) was used. Gel electrophoresis was carried out using 1% SeaKem LE agarose (Lonza, Basel, Switzerland).

For the isolation of antibiotic producing bacteria, three different undefined solid culture media were used: i) Luria-Bertani (LB) agar containing tryptone (1%), NaCl (0.5%), yeast extract (0.5%) and agar (1.5%), pH 7.4 (Sambrook et al., 1989); ii) glycerol yeast extract agar containing glycerol (Sigma-Aldrich)(0.5%), yeast extract (0.2%), K_2_HPO_4_ (0.1%) and agar (1.5%), pH (OpenWetWare, 2013), and iii) starch casein agar containing soluble starch (1%), K_2_HPO_4_ (0.2%), KNO_3_ (0.2%), casein (0.03%)( Sigma-Aldrich), MgSO_4_·7H_2_O (Merck)(0.005%), CaCO_3_ (0.002%), FeSO_4_·7H_2_O (0.001%), agar (1.5%), pH 7.0 (Mohseni et al., 2013). Ultrapure (UP) water, made with a Milli-Q Direct Water Purification System (Merck) was used to make the media. All solutes were purchased from Molar Chemicals Kft., (Halásztelek, Hungary) unless otherwise stated. If fungal contamination was observed, samples were subcultured in the presence of 50 µg/mL cycloheximide (Sigma-Aldrich).

For antimicrobial peptide extraction and analysis, the producer strains were cultured in either of three defined media: i) Mineral Salts medium (MS), containing 17 mM K_2_HPO_4_, 51.5 mM KH_2_PO_4_, 7.6 mM (NH_4_)_2_SO_4_, 0.4 mM MgSO_4_, 1.4 mM Sodium citrate, 2 µM FeCl_2_ and 0.2 % (m/m) glucose (Hall, 1998); ii) rich defined medium (RDM) (Neidhardt et al., 1974) or iii) RDM with Micronutrient Booster solution (Table S2). Amino acids and ribonucleotides were purchased from Sigma-Aldrich (St. Louis, MO, USA).

### Screening of environmental samples for antimicrobial-producing bacteria

About three hundred soil samples were collected from 10-20 cm below ground level, and were screened using the soft-agar overlay assay as follows. A 100 mg sample of the soil was suspended in 1 mL of sterile UP water, vortexed, pelleted at 300 g and the supernatant was streaked out on an agar plate of one of the undefined media described above, in duplicate. The plates were thereafter incubated at 30 and 37 °C for about 3-5 days. After incubation, the colonies were overlaid with pre-cooled (45 °C) soft (0.65%) agar seeded with indicator organisms (*Citrobacter freundii* or *Enterococcus faecalis*). The overlaid plates were incubated at 37 °C overnight, and checked for inhibition zones. Colonies showing zones of inhibition were carefully picked and repeatedly subcultured to obtain a pure culture. The axenic cultures were then subjected to a repeated antimicrobial activity screening using a similar soft-agar overlay assay to confirm the inhibitory activity. Positive colonies were thereafter grown overnight in LB medium and stored in 15% glycerol at -80 °C until further analysis.

### Identification of antimicrobial producers

For the initial, crude identification of antimicrobial-producing isolates, their 16S rRNA gene was PCR-amplified using the primers 27F and 1492R (Table S1)(Hayashi et al., 2004). The amplicon (1500 bp) was purified using the GeneJET PCR Purification kit (Thermo Fisher Scientific, Waltham, MA) according to manufacturer’s instructions. The purified DNA was then deposited for Sanger Sequencing using the primers 27F and 1492R, and the assembled sequencing result was searched in the National Centre for Biotechnology Information (NCBI) database using the BLAST (basic local alignment search tool) algorithm (Altschul et al., 1990).

To refine the phylogenetic analysis of the strain producing the novel AMP (a *Paenibacillus* coded as ISO4), the whole genome sequence of the strain was obtained (see below). The sequence was searched at the NCBI website applying the tBlastn algorithm using the RpoB and GyrA protein sequences of *B. subtilis* as queries to identify the *rpoB* and *gyrA* genes. Next, the concatenated *rpo_gyrA* gene pair of ISO4 was used to query the Refseq reference genomes of the *Paenibacillaceae*, applying nucleotide BLAST search. The obtained top hits were aligned using the MEGA11 software, applying the MUSCLE algorithm. The phylogenetic tree was inferred using the Neighbor-Joining method (Saitou & Nei, 1987). The evolutionary distances were computed using the Tamura-Nei method (Tamura & Nei, 1993).

### Partial purification and concentration of antimicrobial compounds

The AMP-producer bacterial strain was incubated at 37 °C for 24-48 h in 10 mL of one of the defined media described above. The culture was centrifuged at 4500 rpm for 30 min. The supernatant was collected and the active compound was mixed with HPLC grade ethanol (VWR International, Debrecen, Hungary) in 1:4 ratio (v/v) in 15 mL polypropylene tubes. The mixtures were shaken at 2000 rpm for 4 hours at room temperature (Multi Reax orbital shaker, Heildoph, Schwabach, Germany), then centrifuged at 4500 rpm for 30 min. The supernatant was collected in a new 15 mL polypropylene tube and freeze-dried in a lyophilizer (Scanvac Coolsafe, Labogene, Allerød, Denmark). The lyophilized crude extract was suspended in an appropriate volume of HPLC grade water (VWR) to create a 10-fold concentrated solution for activity tests and fractionation.

### Spot-on-lawn assays

The extracts of bacterial cultures were tested for activity against *E. faecalis, Bacillus cereus, Staphylococcus haemolyticus, E. coli, Klebsiella pneumoniae, Pseudomonas aeruginosa and Acinetobacter lwoffii* by a spot-on-lawn assay, as described earlier (Zhao et al., 2020). Briefly, 10 µL of each re-dissolved crude extract was pipetted on the surface of LB-soft (0.65%) agar medium, seeded with one of the bacterial cultures listed earlier. After drying the liquid drops under a sterile hood and overnight incubation of the plates at 37 °C, inhibitory zones were sought in the bacterial lawn.

### HPLC fractionation

Reversed-phase HPLC *separation* was carried out on a Shimadzu LC-20AD system (Shimadzu Corp., Japan) equipped with degasser unit (DGU-20AS), autosampler (SIL-20AC), column oven (CTO-20AC), diode array detector (SPD-M20A) and fraction collector (FRC-10A). Solvent A contained 95% HPLC-grade water (Avantor), 5% HPLC-grade acetonitrile (Avantor) and 0.1% trifluroro-acetic acid (TFA)(Sigma-Aldrich) while solvent B contained 95% acetonitrile, 5% water and 0.1% TFA. Between 10-100 µl of crude extract was loaded on a C18-reversed phase 10 mm x 250 mm x 5 µm (ID x L x particle size) Phenomenex column using a linear gradient of 5-95% B for 50 min, followed by isocratic from 50 to 53 min, then 5% B from 53 to 65 min at a flow rate of 1.5 mL/min for 65 min at 30 °C. About 65 fractions were collected, freeze-dried in a centrifugal evaporator and re-dissolved in HPLC grade water. The fractions were tested for activity against *E. faecalis* by a spot-on-lawn assay, as described above (Zhao et al., 2020). The active fractions from the first HPLC run were subjected to a second or third rounds of HPLC separations on a 2.1 mm x 50 mm x 2.5 µm (ID x L x particle size) Waters BEH C18-reversed phase column. The pure biologically active fractions were then subjected to Mass Spectrometry (MS).

### LC-MS analysis

LC-MS/MS data were acquired using a NanoAcquity UPLC (Waters) online coupled to an Orbitrap Elite (Thermo Scientific) mass spectrometer. 5 µl of sample was injected onto a trapping column (Waters m-Class Symmetry, part #:186007496, 0.180 mm x 20 mm x 5 µm (ID x L x particle size)) using 1% solvent B (solvent A: 0.1% FA/water, solvent B: 0.1% FA/ACN, FA: formic acid, ACN: acetonitrile) at a flow rate of 5 µl/min for 4 min then transferred onto the separating column (Waters nanoAcquity UPLC BEH130 C18, part #:186003544, 0.075 mm x 200 mm x 1.7 µm (ID x L x particle size)) using gradient elution: 5% B from 0 to 5 min, 5 to 10% B from 5 to 7 min, 10-50% B from 7 to 32 min, 50 to 90% B from 32 to 33 min, 90% B from 33 to 38 min, 90 to 5% B from 38 to 40 min followed by equilibration of the column with 5% B for 20 min. Flow rate was 250 nl/min and the column was thermally stabilized at 45 °C. Effluent of the column was directly introduced into the mass spectrometer using electrospray ionization followed by data dependent analysis acquiring high-resolution MS followed by collecting CID (collision induced dissociation) and HCD (higher energy collision dissociation) MS/MS data of the most abundant multiply charged precursor ions. MS data were collected in the m/z range of 380-1400. Both MS and MS/MS data were acquired using the Orbitrap analyzer at a resolution of 60000 and 15000 for MS and MS/MS data, respectively. Normalized collision energy was set at 32 and 35 for HCD and CID activation, respectively. Dynamic exclusion was enabled to preclude repeated fragmentation of precursor ions already analyzed (exclusion time: 10 s, precursor mass accuracy: ±10ppm).

Comparison of the synthetic polymyxins and the active fraction of ISO4 followed the above setup except that a different gradient elution program was used: 5% B from 0 to 5 min, 5 to 30% B from 5 to 10 min, 30-50% B from 10 to 30 min, 50 to 90% B from 30 to 33 min, 90% B from 33 to 38 min, 90 to 5% B from 38 to 40 min followed by equilibration of the column with 5% B for 20 min.

The active fraction of the ISO4 isolate was also analyzed by an Evosep One-Orbitrap Fusion Lumos Tribrid setup. Samples were analyzed using the “30SPD” built-in 44-min gradient of the Evosep system with identical MS settings described above for the Orbitrap Elite MS except that MS and MS/MS data were acquired at a resolution of 120000 and 30000, respectively and HCD activation was performed at NCE: 30.

### Whole genome sequencing and identification of biosynthetic gene clusters

The chosen AMP-producer strains were fully grown in LB medium overnight. Genomic DNA was extracted using the NucleoSpin Microbial DNA Mini kit (Macherey Nagel, Düren, Germany) according to the manufacturer’s instructions. Whole genome sequencing was carried out by Delta Bio 2000 Ltd. (Szeged, Hungary). The obtained DNA contigs were screened for biosynthetic gene clusters responsible for antimicrobial peptide production using the antiSMASH web server (Blin et al., 2023).

### Electroporation of bacterial strains

Electrocompetent *E. coli* cells were prepared and transformed using regular protocols (Shukla et al., 2022). For electroporation of *Paenibacillus* strains, the method of Li *et al* was used. Briefly, a *Paenibacillus* starter culture grown in LBS medium (LB + 0.5 M sorbitol) overnight was diluted 100-fold into 20 mL of fresh LBS and cultured at 37 °C until an optical density (OD590) of 0.8-1.0. After 10 min on ice, the culture was centrifuged at 6300 rpm for 15 min at 4 °C. The pellet was collected and washed four times with electroporation buffer (0.5 M sorbitol, 0.5 M mannitol and 10% glycerol), finally resuspending in 100 µL of the same buffer. Electroporation was carried out in 1 mm electroporation cuvettes (Cell Projects Ltd, Kent, UK) with a Micropulser electroporator (BioRad, Hercules, CA, USA) set at 2000 V. After electroporation, cells were collected in 1 mL of recovery medium (LB + 0.5 M sorbitol + 0.38 M mannitol) and shaken at 220 rpm at 30 °C for 3-5 hours prior to plating on the appropriate antibiotic containing plates. Colonies were PCR-screened after incubation at 30 °C for 24-48 hours (Z. Li et al., 2020).

### Genome editing of Paenibacillus

Plasmid pJOE26_16 was transformed into *Paenibacillus* strain ISO4 using the above protocol. Verified transformants, selected on LB + Km-plates at 30 °C, were restreaked on LB + Km plates and incubated at 37 °C. Plasmid co-integrants were identified using colony PCR: plasmid cointegration through left homology was indicated by a positive 26_16checkA + 26_16checkC PCR (700 bp product). Plasmid co-integration through right homology was indicated by a positive 26_16checkB + 26_16checkD PCR (640 bp product) (See Table S1 for primer sequences). A verified plasmid co-integrant clone was picked and grown in liquid LB + 0.2% mannose solution to induce CRISPR/Cas cleavage, diluted and plated on LB + mannose plates to obtain individual colonies. Double cross-over, i.e. complete deletion of the targeted chromosomal segment and elimination of the integrated plasmid was declared if the PCR with primers 26_16checkA + 26_16checkD displayed the correct length (deletion mutant: 1140 bp, WT: 3000 bp).

### Peptide synthesis

Four selected peptide sequences were synthesized by Genscript (Rijswijk, Netherlands), in 5 mg quantities each. The peptides were resuspended in HPLC-grade water (VWR), diluted to 0.5 µg/mL using 0.1% formic acid and were analyzed using the LC/MS setup as described above, for verification of their mass and comparison of their retention times and fragmentation spectra to those of the peptides present in the purified bacterial extract.

### Recording the antimicrobial spectrum

The test strains included 10 aerobic Gram-negative non-fastidious, 2 aerobic fastidious, 8 Gram-positive and 1 anaerobic organism (Table 4). For determination of the MICs of the synthetic polymyxin variants, the broth-microdilution method was applied as recommended by international clinical standardization bodies (EUCAST, CLSI). The following media were applied: cation-supplemented Müller-Hinton broth (CAMH) for non-fastidious bacteria, CAMH supplemented with 5% lysed horse blood, 20 μg/mL β-NAD for fastidious aerobic bacteria and supplemented Brucella broth (5% lysed horse blood, 5 μg/mL hemin and 1 μg/mL vitamin K_1_) for the anaerobic bacterium. To describe the process briefly, appropriate concentrations of the compounds were 2-fold diluted serially in sterile ultrapure water (100 μL) and ca. 10^5^ test organism cells in 100 μL of 2x concentrated broth were added that were prepared by addition of 1/100^th^ of a 0.5 McFarland cell suspension in PBS. The working antibiotic concentration ranged from 128 μg/mL to 0.25 μg/mL with a blank control. Aerobic isolates were incubated at 37 °C in a CO_2_ thermostat for 24 h, the anaerobic isolate was incubated anaerobically in an anaerobic cabinet (85% N_2_, 10% H_2_ and 5% CO_2_, Whitley A45 anaerobic workstation, Bentley, UK) at 37 °C for 48 h, after which visible growth was recorded.

### Analysis of toxicity

Three cell lines were used for toxicity measurements. HepG2 (ATCC, Manassas, VA, USA) is a human hepatoblastoma carcinoma cell line of epithelium morphology. HEK293 (ATCC) is a human embryonic kidney cell line. HRPTEpC (Sigma-Aldrich, St. Louis, MO, USA) is a human renal proximal tubular epithelial cell line. Cytotoxicity of the peptides was checked by XTT colorimetric reaction, which measures cell metabolism, proportional with the living cell numbers. The HepG2 and HEK293 cell lines were plated at 2x10^4^ cells/well, while the HRPTEpC cells at 10^4^ cells/well in DMEM-F12 medium with 10% FCS (Gibco, Merck KGaA, Darmstadt, Germany) in flat bottom 96 well tissue culture plates (Orange Scientific, Braine-l’Alleud, Belgium). After overnight incubation the peptides were administered in fresh medium at doubling dilutions starting from 100 µg/ml. The XTT assays were performed according to the instructions of the manufacturer (CyQUANT XTT Cell Viability Assay, Thermo Fisher Scientific Inc., Waltham, MA, USA). The IC_50_ concentrations, i.e, the minimal peptide concentrations resulting in 50% decrease of XTT specific absorbance (OD_450_ – OD_690_), were determined after 24 h.

## Results

### Functional screens of environmental samples

The core research process of this work was to conduct environmental screens with the aim of discovering novel AMPs. The environmental samples originated from various soils and seawater. For simplicity, we refer to mainland soils as “soil”, to seabed soils as “lagoon”, and to seawater as “sea” when referring to the source of sampling (**Table 1**). A total of cca. 300 samples were collected, processed and screened using classical agar overlay assays, as described above. To increase the odds of observing hits (i.e. zones of inhibition), both Gram-negative and Gram-positive strains (*C. freundii* and *E. faecalis,* respectively) were used as target strains in parallel. Positive colonies were recovered and subcultured to obtain axenic cultures. The supernatants of these cultures were used for activity verification using spot-on-lawn assays. Sanger sequencing of the amplified 16S rRNA gene of each strain was used for their crude identification and dereplication. A total of 29 bacterial strains belonging to six genera (**Table 1**) were isolated and identified this way. Notably, one of our isolates was an anaerobic antimicrobial-producing bacterium (*Clostridium beijerinckii*).

**Table 1.** List of antimicrobial-producing bacteria isolated in this study. The sample that this study focuses the most on is in bold.

| Sample number | Source | Closest related species | Percent identity (%) | Accession No. of closest related sp. |
| --- | --- | --- | --- | --- |
| 1 | soil | *Bacillus cereus* | 98.09 | OP778612.1 |
| 2 | lagoon | *Bacillus sp.* | 97.7 | OL468372.1 |
| 3 | soil | *Brevibacillus parabrevis* | 97.04 | CP118544.1 |
| 4 | soil | *Brevibacillus brevis* | 98.34 | LR134338.1 |
| 5 | soil | *Brevibacillus spp* | 96.85 | MN577296.1 |
| 6 | soil | *Brevibacillus laterosporus* | 96.99 | DQ371289.2 |
| 7 | soil | *Brevibacillus sp.* | 97.71 | KT363757.1 |
| 8 | soil | *Brevibacillus sp.* | 98.02 | HF545882.1 |
| 9 | lagoon | *Clostridium beijerinckii* | 97.63 | KX269863.1 |
| 10 | lagoon | *Paenibacillus alvei* | 96.13 | KF793042.1 |
| 11 | soil | *Paenibacillus alvei* | 98.14 | JF830167.1 |
| 12 | soil | *Paenibacillus alvei* | 97.41 | MT498465.1 |
| 13 | soil | *Paenibacillus alvei* | 97.75 | JF701948.1 |
| **14** | **soil** | ***Paenibacillus sp. (ISO4)*** | **98.12** | **CP121215.1** |
| 15 | soil | *Paenibacillus dendritiformis* | 98.83 | KF850540.1 |
| 16 | soil | *Paenibacillus dendritiformis* | 97.88 | OX216966.1 |
| 17 | soil | *Paenibacillus elgii* | 97.87 | MZ026448.1 |
| 18 | lagoon | *Paenibacillus elgii* | 97.44 | KT719740.1 |
| 19 | lagoon | *Paenibacillus polymyxa* | 97.66 | EU982535.1 |
| 20 | lagoon | *Pseudomonas mendocina* | 97.63 | CP002620.1 |
| 21 | lagoon | *Pseudomonas mendocina* | 97.77 | LC107430.1 |
| 22 | lagoon | *Pseudomonas aeruginosa* | 100 | MT785904.1 |
| 23 | soil | *Streptomyces lilaceus* | 97.56 | OR434662.1 |
| 24 | soil | *Streptomyces rimosus* | 98.7 | PP954962.1 |
| 25 | sea | *Streptomyces vinaceusdrappus* | 98.58 | KC747480.1 |
| 26 | sea | *Streptomyces globosus* | 97.5 | MN538259.1 |
| 27 | sea | *Streptomyces chryseus* | 98.02 | EU841613.1 |
| 28 | sea | *Streptomyces thermolilacinus* | 96.34 | OP271846.1 |
| 29 | sea | *Streptomyces sp.* | 98.1 | MW695205.1 |

After having tested various extraction methods and organic solvents, we extracted the antimicrobial compounds from the supernatants of the axenic cultures. The antimicrobial spectra of these extracts were qualitatively assessed using spot-on-lawn assays targeting 3 Gram-positive and 4 Gram-negative bacterial strains (**Table 2**). In the downstream analysis, we focused on the genus *Paenibacillus* for two reasons: i) this genus had the highest number of representatives (10) among the positive strains and ii) our preliminary result showed that isolates belonging to genus *Paenibacillus* (and *Brevibacillus)* were cultivable and produced their antimicrobial compound in defined media, unlike *Streptomyces* which were not producing antimicrobial compounds in such media. We expected analytical procedures to produce more reproducible results with lower backgrounds when using defined media for cultivating the strains from which the crude extracts are prepared, since there is no lot-to-lot variation and the unknown composition of undefined growth media is avoided.

**Table 2.**
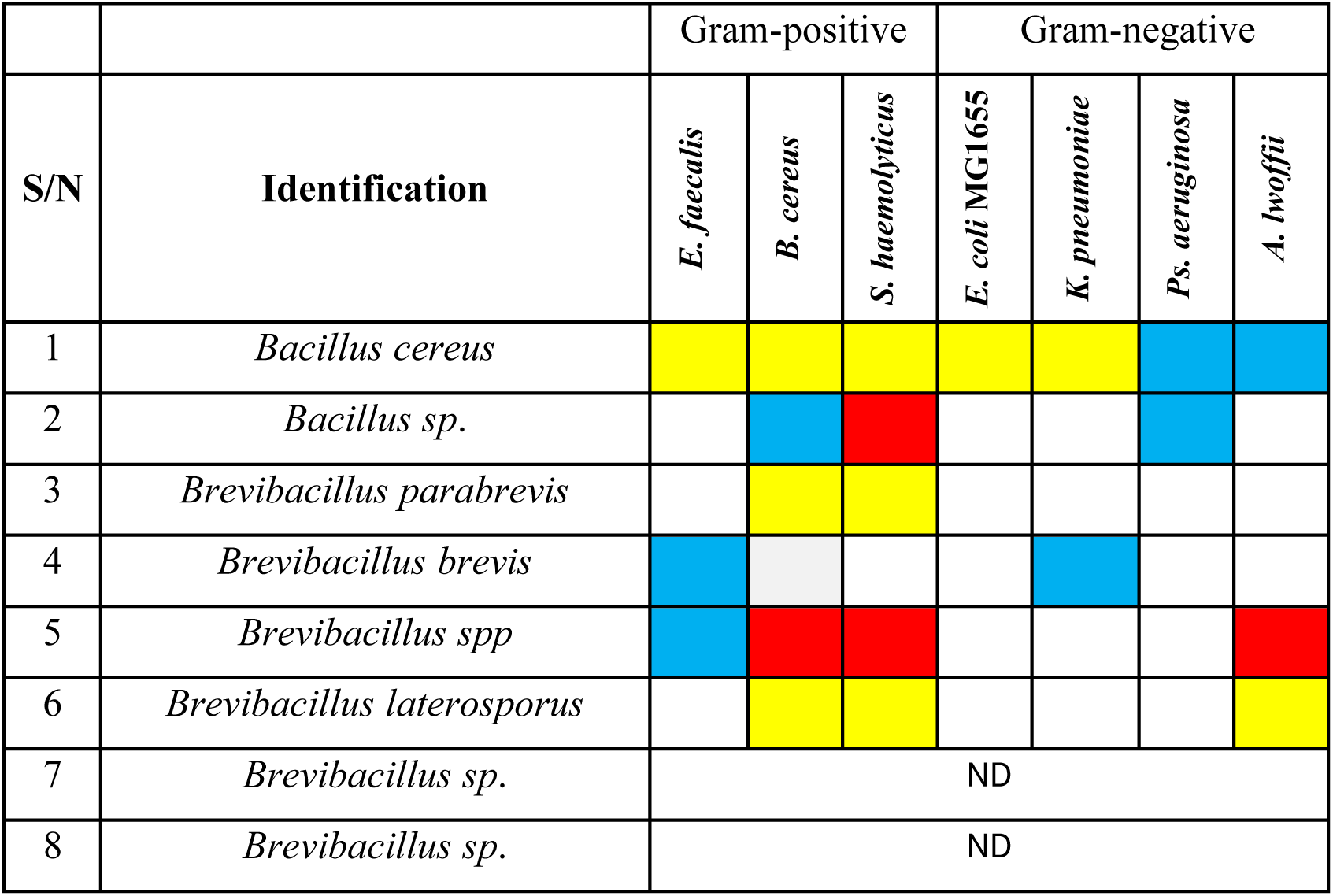
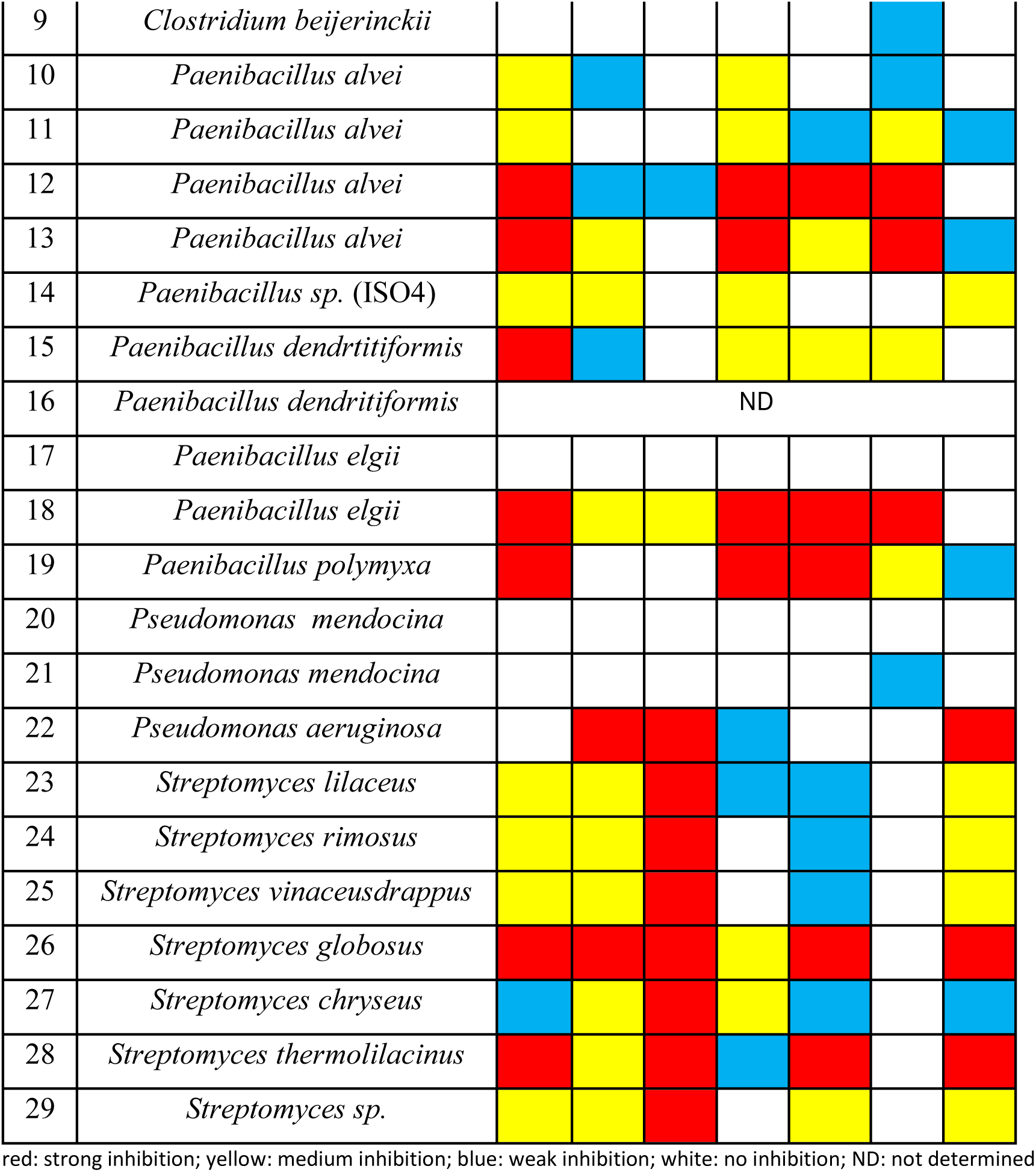
Activity of supernatant extracts on a panel of target bacteria.

The crude extracts of the analyzed *Paenibacilli* underwent reversed phase HPLC separation on a semi-preparative C18 column. The fractions were spotted on *E. faecalis* lawns, and active fractions were re-fractionated in a second and third dimension HPLC run using a C18 column with different separation properties. Spot-on-lawn assays were repeatedly used to define active fractions (**Figure S1**). Fractions that showed inhibitory activity along with neighboring inactive fractions (as negative controls) were submitted for LC-MS/MS analysis. MS/MS spectra of the dominant, multiply charged precursor ions within the LC-MS profiles were subjected to manual *de novo* sequencing.

Fragmentation patterns were interpreted using known *Paenibacillus* AMP sequences as templates, which revealed that all active peptides of isolate ISO4 (sample 14) belonged to the polymyxin family. A number of other paenibacilli isolated by our lab produced known polymyxins, as deduced from their mass spectrometric data (MS and MS/MS spectra) (**Table 3**). Sample 14 (ISO4) however secreted a polymyxin with a unique molecular weight, represented by the dominant ion of m/z 389.936(3+).

**Table 3.**
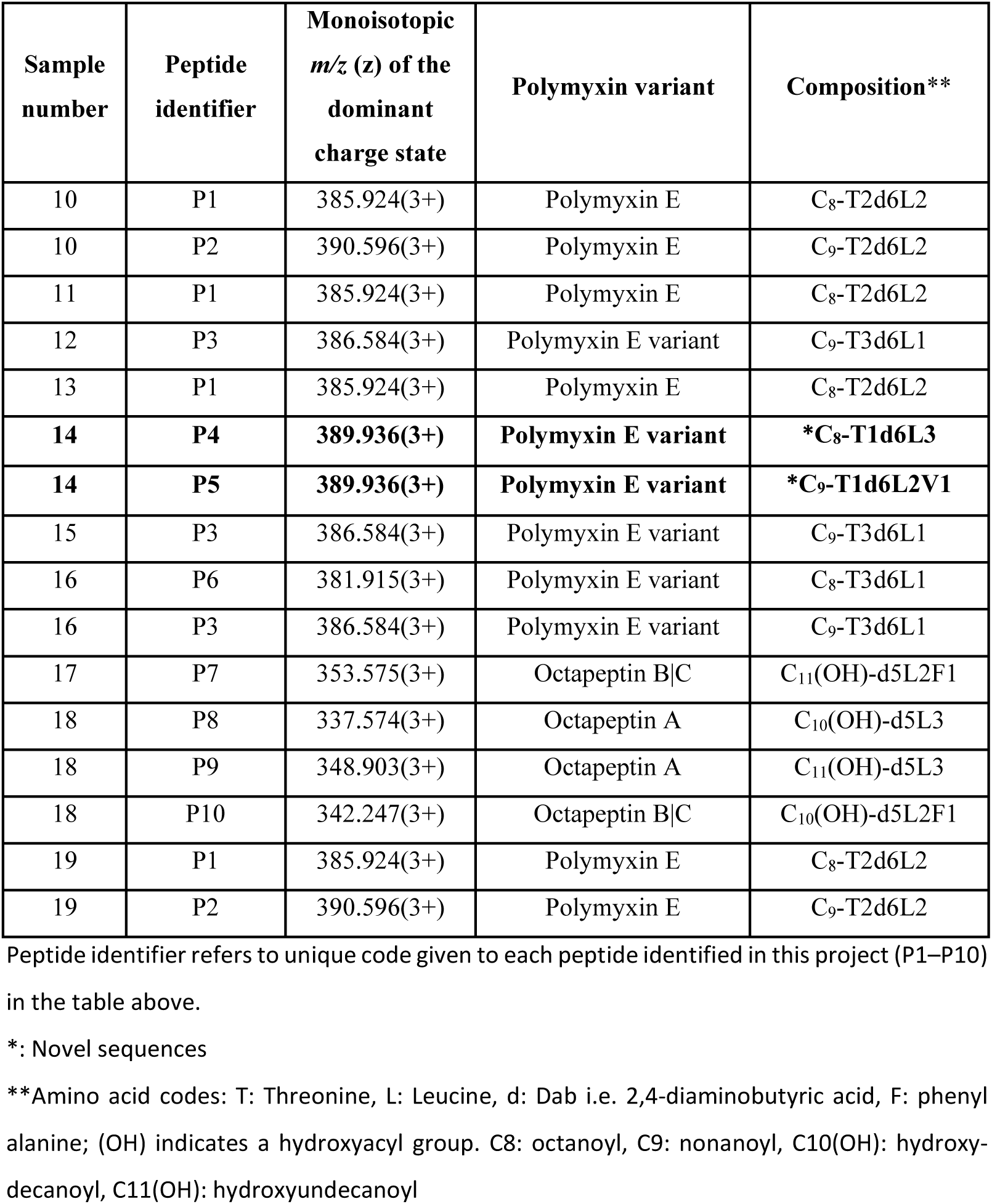
List of polymyxin variants identified by mass spectrometry in *Paenibacillus* strains.

Monitoring for the presence of this new polymyxin variant in subsequent ISO4 extracts revealed that this ion was attributable to two, partly coeluting peptides of identical molecular weight: one carrying three Leu/Ile residues (**P4 of Table 3**), the other carrying one Val and two Leu/Ile residues within the cyclic ring (**P5 of Table 3)**. Manual inspection of the ion trap CID spectra of the doubly charged peptides revealed that the earlier eluting P5 (**Figure 1, top panel**) carries a nine-carbon N-terminal acyl group, while the later eluting P4 is modified with an eight-carbon acyl group (**Figure 1, bottom panel**) as demonstrated by the abundant b_1_ ions of m/z 241 and 227, respectively. The 14-Da mass difference between the acyl groups is compensated by a Leu -> Val substitution in the earlier eluting peptide as demonstrated by a set of fragment ions (connected by dashed arrows in Figure 1) such as m/z 313 and 327 representing b-type fragment ions of Leu-Val-Dab and Leu-Leu-Dab compositions in P5 and P4 peptides, respectively. Furthermore, the m/z 313 fragment ion also proves that the Val is directly connected to one of the Leu residues. Thus, considering the conserved amino acid positions in natural polymyxins we presume that the Val must be in position 7. Note that different isomers of the peptide building blocks cannot be distinguished by the current LC-MS/MS setup, e.g. we cannot tell whether the N-terminal acyl groups and amino acid side chains are linear or branched, or the amino acids are of D- or L relative configuration. For simplicity we refer to all building blocks of a residual mass of 113 as Leu keeping in mind that these might also be Ile or even Nle (norleucine). Furthermore, although the fragment at m/z 213.160 (**Figure 1, upper panel**) confirms a direct Val-Leu linkage, the exact position of Val within the cyclic core cannot be unambiguously assigned based on MS/MS data alone. Due to the high number of repetitive residues (four Dab and two Leu) and the characteristic multiple ring-opening pathways of cyclic peptides, alternative cleavage events generate identical isomeric fragment ions. To narrow down the theoretical structural possibilities, we relied on established literature, which defines positions A6, A7, and A10 as the highly variable regions in known polymyxin peptides. Guided by this structural consensus and the direct evidence for the Val-Leu linkage, we assigned these two residues to the A6–A7 positions, although the sequence remains indistinguishable between the Val-Leu and Leu-Val permutations.

**Figure 1.**
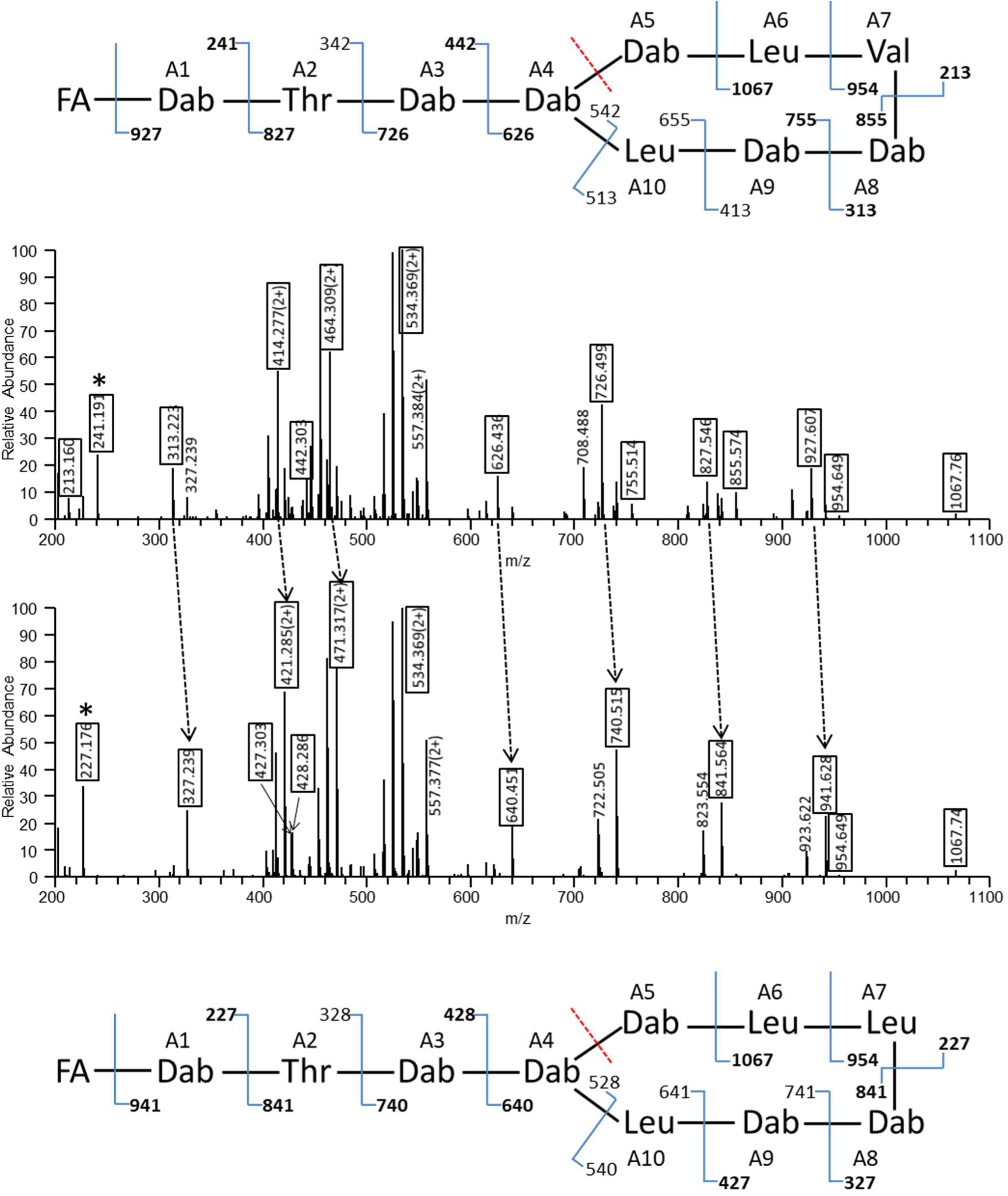
CID spectra of the doubly charged precursor ions at m/z 584.402, which correspond to the novel polymyxin peptides identified in ISO4. The earlier eluting P5 peptide (top panel) is N-capped with a C9-fatty acyl (FA) group and contains a Val residue in the ring, while the later eluting P4 peptide (bottom panel) is N-capped with a C8-acyl group and features three Leu’s in the cyclic region. Potential peptide bond fragment ion series starting from the N-terminus or A5|A6 cleavage site are listed along with the structures deciphered and the ones observed are highlighted in the spectra. b_1_ ions, indicating differential acylation are labeled with asterisks, ion pairs demonstrating the Val-Leu ’swap’ are connected by dashed arrows. Dab: 2,4-diaminobutyric acid.

### Genome scale analysis of the Paenibacillus ISO4 strain

After detection of the potentially novel polymyxin variants in the active fractions of the ISO4 extract, we proceeded to identify the genes responsible for the production of the polymyxin compounds. First, we isolated genomic DNA from the strain and performed whole genome sequencing. The sequence was deposited to the antiSMASH webserver (Blin et al., 2023) to identify biosynthetic gene clusters (BGC) responsible for secondary metabolite synthesis. The antiSMASH analysis predicted 15 clusters to encode non-ribosomal peptide synthases (NRPS), polyketide synthases (PKS), and other assembly line biosynthetic gene clusters **(Figure S2)**. Most notably, a complete NRPS-type BGC, sufficient for colistin (polymyxin E) synthesis was annotated on contig 26. The annotated BGC was 41.1kb long and consisted of five open reading frames (ORFs). Three of these, ctg26_11, ctg26_13 and ctg26_16 corresponded to the previously described 3-core biosynthetic polymyxin synthetase genes *pmxA, pmxB* and *pmxE,* respectively. Two ORFs (ctg26_14 and ctg26_15) corresponded to transport-related genes *pmxC* and *pmxD,* respectively (**Figure 2**). As expected, the polymyxin synthetase genes consist of series of modules responsible for the incorporation of amino acid residues corresponding to their sequence within the AMP. Each module’s condensation, adenylation and thiolation domains were readily identified by the antiSMASH platform. The adenylation domains are responsible for selecting the amino acid residue to be incorporated, and antiSMASH also provided the specificity of each such domain. Relying on earlier studies (Choi et al., 2009), we used the order of adenylation domains lined up in pmxE, pmxA and pmxB (in this order) to reconstruct the N-to-C sequence of the polymyxin peptides. The presence of epimerase domains in certain modules (namely 3 and 6) aided us in predicting which amino acids are epimerized to their D-enantiomer (**Figure 2**). Based on this, we refined the MS/MS-based prediction of P4 to be: **octanoyl-Dab-Thr-dDab-cy(Dab-Dab-dLeu-Ile-Dab-Dab-Leu).** The prediction of A7 was ambiguous or missing during repeated antiSMASH analyses, we therefore included the predictions of PARAS (Terlouw et al., 2026) to further refine the output. PARAS predicted A7 to be Ile (probability score 0.465), Leu (0.375) or Val (0.033). As for A6 and A10, PARAS predicted Leu with a probability score of 0.962, and did not include Val among the top ten predicted residues. We therefore concluded that Val most probably lies at position A7, and the sequence of P5 is therefore most likely **nonanoyl-Dab-Thr-dDab-cy(Dab-Dab-dLeu-Val-Dab-Dab-Leu).**

**Figure 2.**
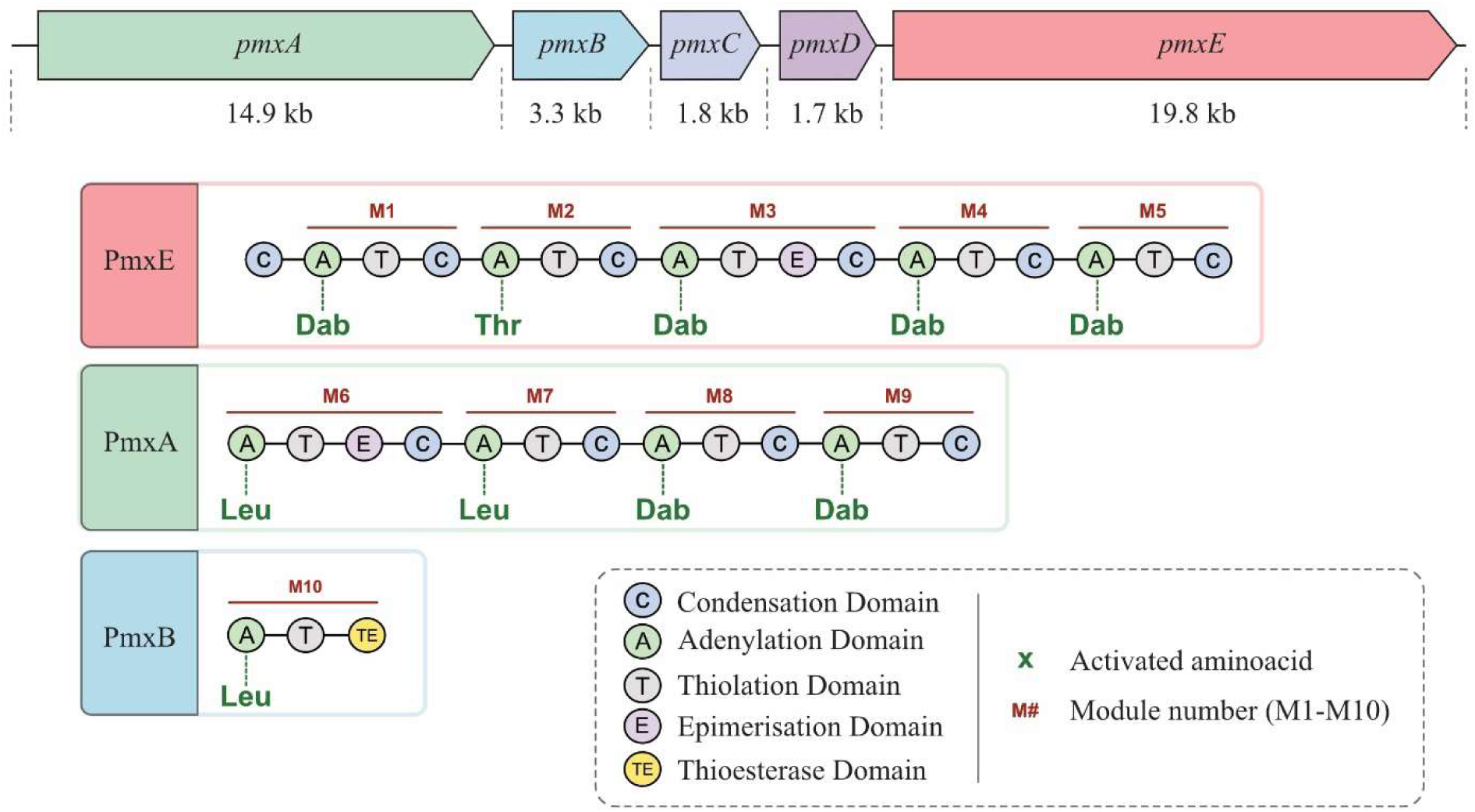
Top: The identified gene cluster for polymyxin biosynthesis in *Paenibacillus* strain ISO4. Middle: the order of NRPS modules responsible for polymyxin synthesis. The specificities of the adenylation domains are indicated. Bottom: the color coding of the domains present in the NRPS modules. The image was made using BioRender.

The availability of the whole genome sequence of *Paenibacillus* strain ISO4 allowed us to refine the phylogenetic mapping of the strain using a more precise approach (see Methods). Based on the sequences of the *rpoB* and *gyrA* genes, strain ISO4 was found to most closely resemble *Paenibacillus barciconensis* strain KACC11450 within the RefSeq database (**Figure S3**).

### Validation of the polymyxin BGC via genome editing

The antiSMASH webserver identified only a single BGC as an NRPS capable of polymyxin synthesis, experimental evidence was therefore needed to investigate whether it was responsible for the production of both novel polymyxins. To obtain this evidence, we engineered a 1.8 kbp deletion in the ctg26_16 (*pmxE*) gene of strain ISO4. The deletion was obtained using a hybrid approach described earlier (**Figure S4** in the Supplement)(Altenbuchner, 2016): first, suicide plasmid pJOE26_16, carrying two genomic fragments was transformed into *Paenibacillus* strain ISO4. Plasmid cointegration into the genome was selected by culturing transformants in the presence of Km at increased temperatures. After PCR-verification of cointegration, the CRISPR/Cas-mediated cleavage of the gene to be deleted was induced, thereby selecting for a molecular resolution that removes the target gene and the inserted cassette altogether. The scarless deletion within the ctg26_16 gene was verified by PCR-amplification of the genomic region, followed by Sanger sequencing of the PCR fragment.

The obtained deletion mutant strain ISO4Δ26_16 went through four levels of phenotypic testing. First, in a soft agar overlay assay, we observed no inhibitory zones around the ISO4Δ26_16 colony (**Figure S5**). Second, extraction and HPLC separation of the spent media originating from the cultured deletion strain was carried out, followed by spot-on-lawn assays of the obtained fractions. Neither of the fractions displayed antibacterial activity (**Figure 3**.).

**Figure 3.**
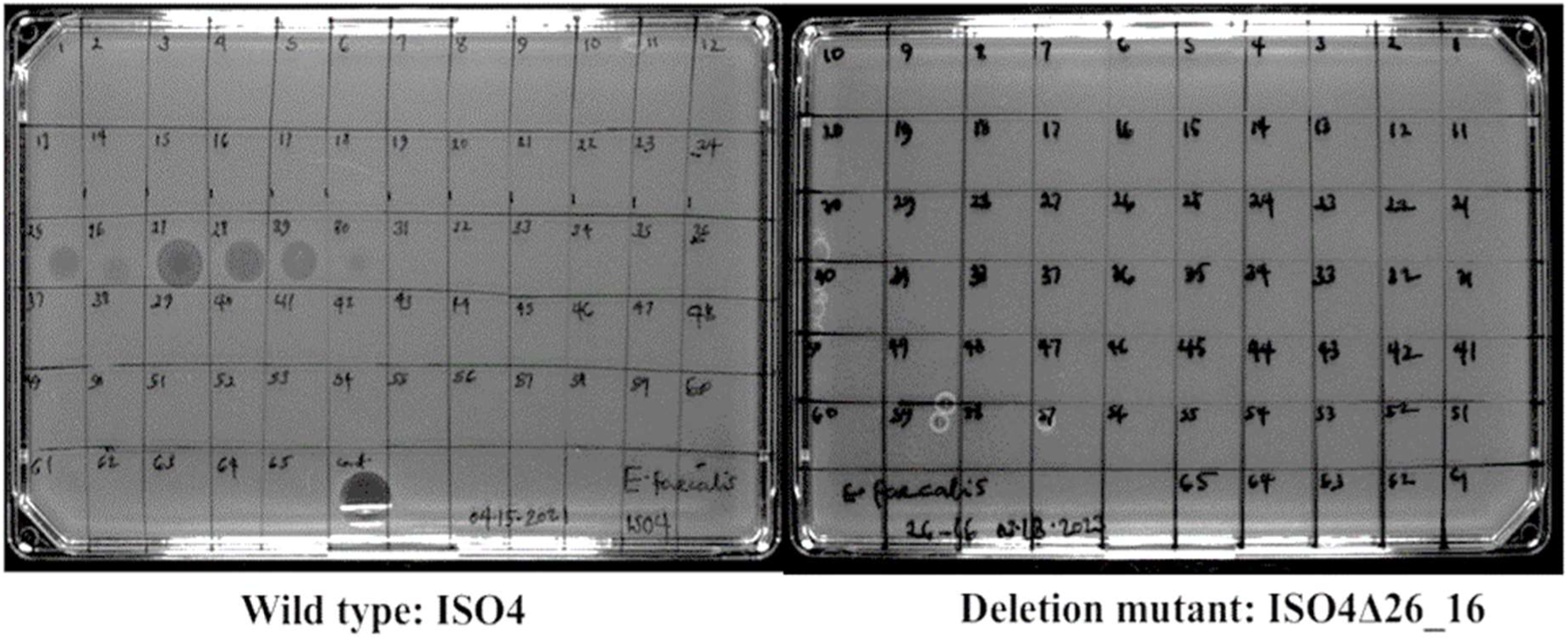
Comparison of antibacterial activities of the 65 HPLC fractions from wild-type (ISO4, WT: left figure) and deletion mutant (ISO4Δ26_16: right figure) of *P. barciconensis* (ISO4) against *E. faecalis*. The activity seen on the bottom of the left panel is 5 µL of the injected crude extract spotted directly as control.

Third, the active HPLC fraction (fr. 27) of the wild type ISO4 strain extract and the corresponding inactive fraction of the ISO4Δ26_16 strain extract underwent LC-MS/MS analysis. In the base peak intensity (BPI) trace of the MS output, no peaks corresponding to polymyxins could be seen in the ISO4Δ26_16 strain (**Figure 4**). And finally, the peaks corresponding to the new polymyxin (*m/z* 584.4(2+) and 389.94(3+)) were under limit of detection in the extracted ion chromatogram (EIC) of fraction number 27 of the mutant ISO4Δ26_16 (**Figure 5**). These phenotypic changes confirmed that the NRPS cluster identified on the ctg26 contig was the only BGC responsible for the synthesis of the novel polymyxins.

**Figure 4.**
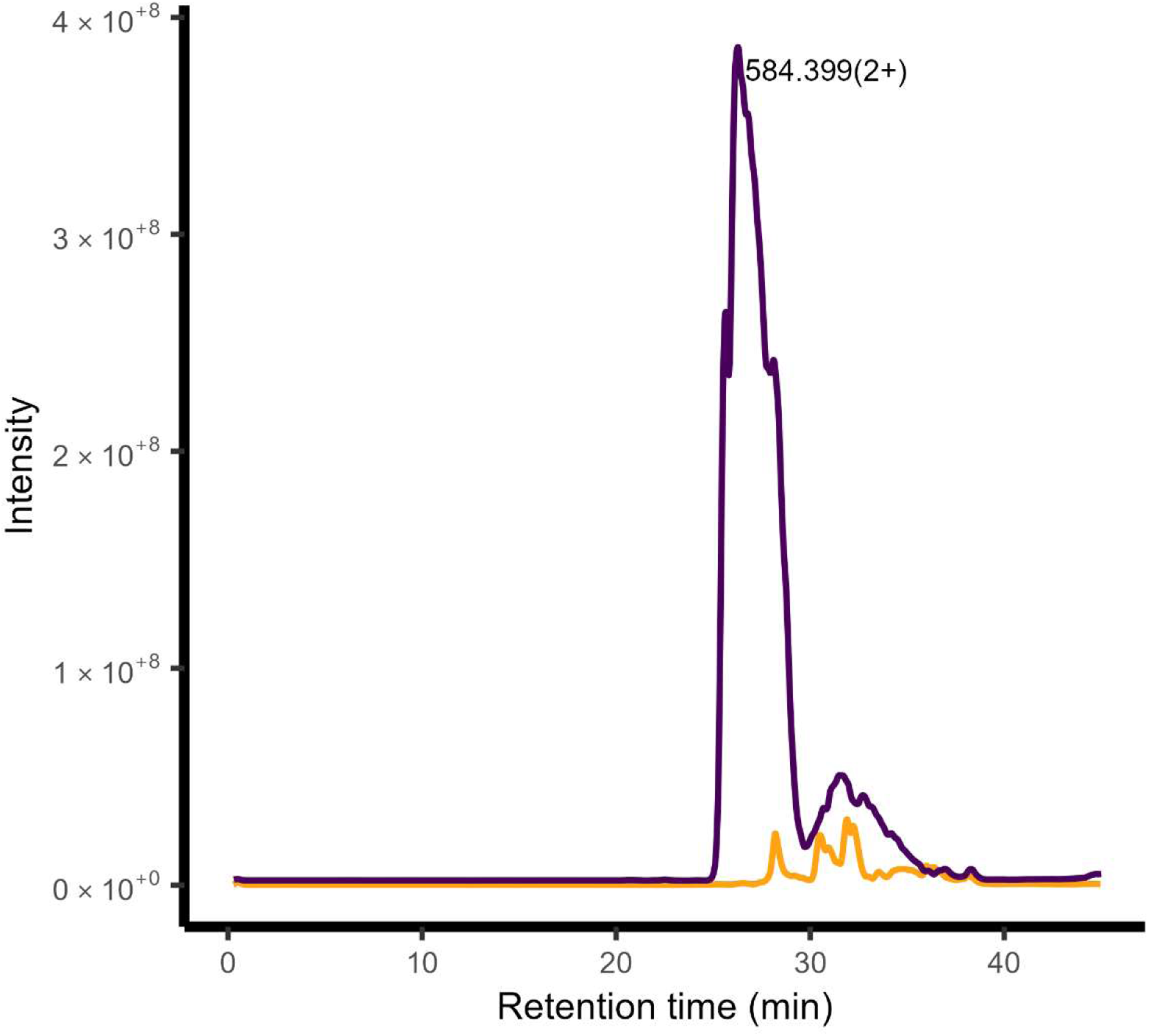
Effect of 1.8 kb deletion in *pmxE* gene of. *P. barciconensis* (ISO4) on the presence of polymyxin within the fractionated extract. Base peak intensity (BPI) profiles of polymyxin *m/z* 584.399(2+) in HPLC fraction number 27 obtained from *P. barciconensis* ISO4 (purple line) or deletion mutant ISO4Δ26_16 (orange line) strain.

**Figure 5.**
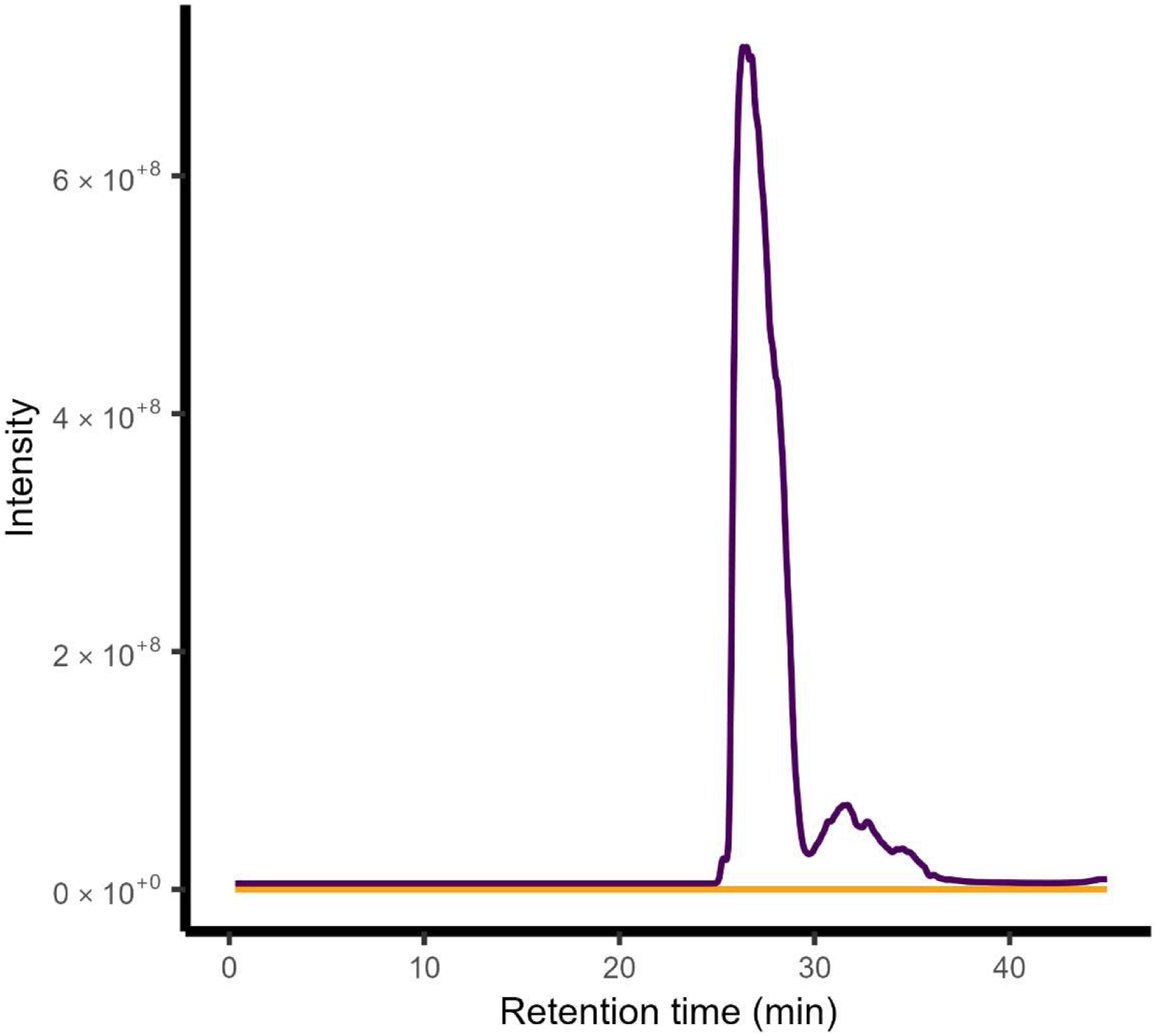
Comparison of MS-extracted ion chromatograms (*m/z* 389.936 and 584.400 representing the triply and doubly charged precursor ions of the new polymyxin, respectively) from fraction number 27 of wild-type (purple line) and deletion mutant Δ26_16 (orange line) of *P. barciconensis* (ISO4).

### Synthesis and quantitative analysis of novel polymyxin variants

Due to the inherent inability of MS/MS to differentiate between isomeric forms (such as Leu/Ile structural isomers or D/L stereoisomers) at specific positions, combined with the redundant fragment ions generated by multiple ring-opening pathways of the cyclic core, unambiguous de novo sequencing was precluded. Therefore, further strategies were needed to take our polymyxin-discovery project to a conclusive stage. Ideally, heteronuclear 2D-NMR analysis could complete the identification process of the novel AMPs. However, this approach would require milligram quantities of the peptides in high purity, both of which were technically out of our reach. We therefore decided not to pursue the exact identification of the natural peptides, but instead, use the available chemical information to design multiple novel, previously undiscovered polymyxins. The peptides to be synthesized were chosen to be novel active AMPs closely related (or identical) to the ones extracted from the natural isolates. In this respect, four synthetic polymyxins were designed. Two of these, marked C81 and C82, harbored a C_8_-acyl chain at the N-terminus, while the other two (C91 and C93) carried a C_9_-acyl chain (**Figure 6**). All acyl chains lacked branching of any kind. C81 had D-Leu, Ile and Leu residues in positions 6, 7 and 10, respectively. C82 carried D-Leu, Leu and Leu residues in these positions. Both C91 and C93 carried D-Leu, Val and Leu residues in these positions, the only difference lay in their N-terminal tripeptide sequences: C91 had the conventional Dab, D-Thr, D-Dab, while C93 comprised the unconventional Dab, D-Dab, D-Thr at positions 1, 2 and 3, respectively. These two were synthesized with the aim of testing the effect of the amino-acid swap of A2 and A3 on the MS/MS spectrum and the activity (explained in the Supplementary discussion). Total synthesis of the peptides was outsourced to a peptide synthesis company. Upon arrival, our in-house LC-MS/MS analysis verified the correct composition and the high (>95%) purity of all four synthetic polymyxins (data not shown). The retention times of the four synthetic polymyxins were distinct from those of the two natural variants found in the active fractions of *Paenibacillus* strain ISO4 (**Figure S6**).

**Figure 6.**
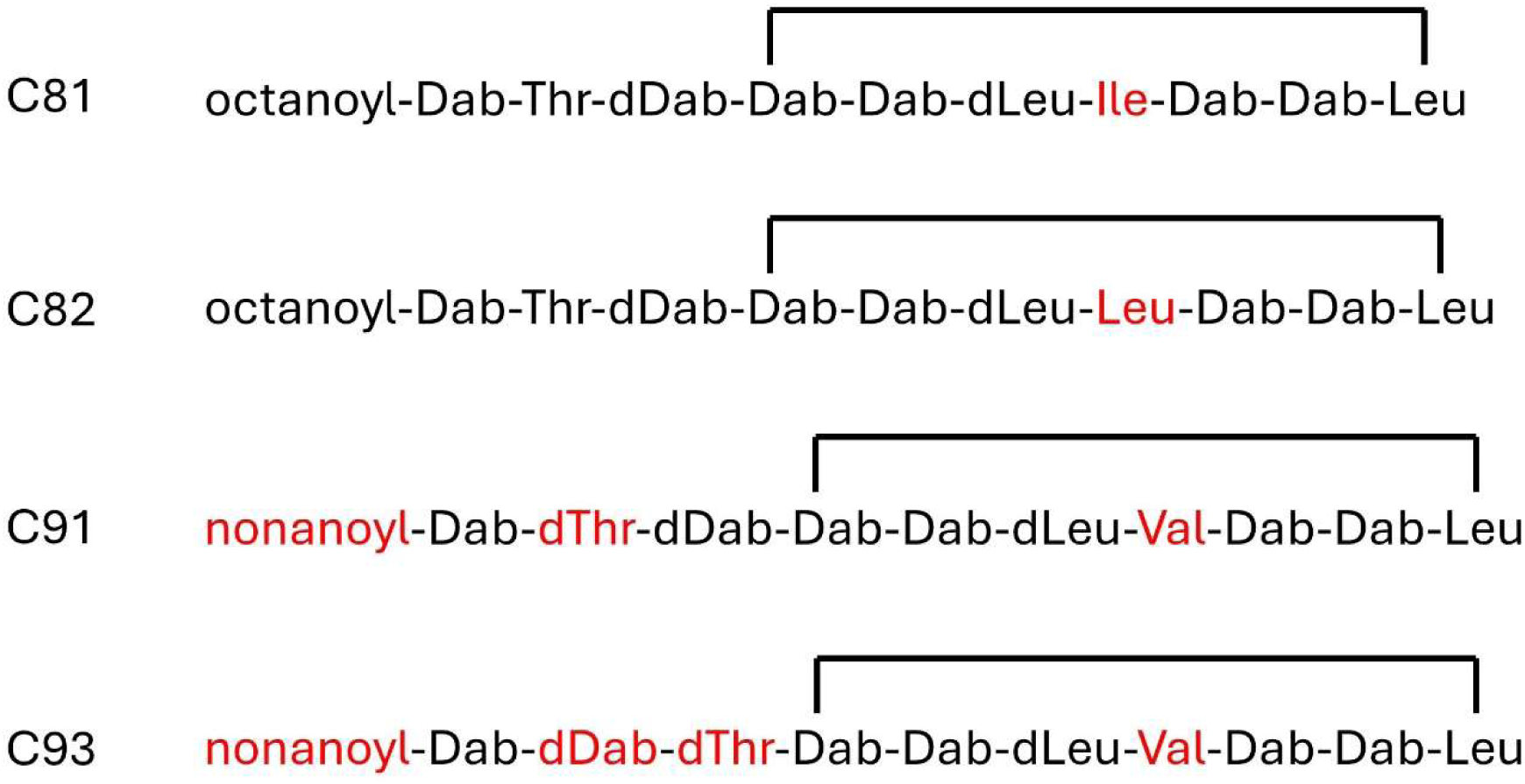
Schematic representation of the four synthetic polymyxins. The label of each polymyxin is depicted on the left. Thick lines within the structural formulas mark cyclisation (the C-terminal carboxyl group making an isopeptide bond with the side chain amine of the Dab at position 4). Variations within the four sequences are highlighted in red.

**Figure 7.**
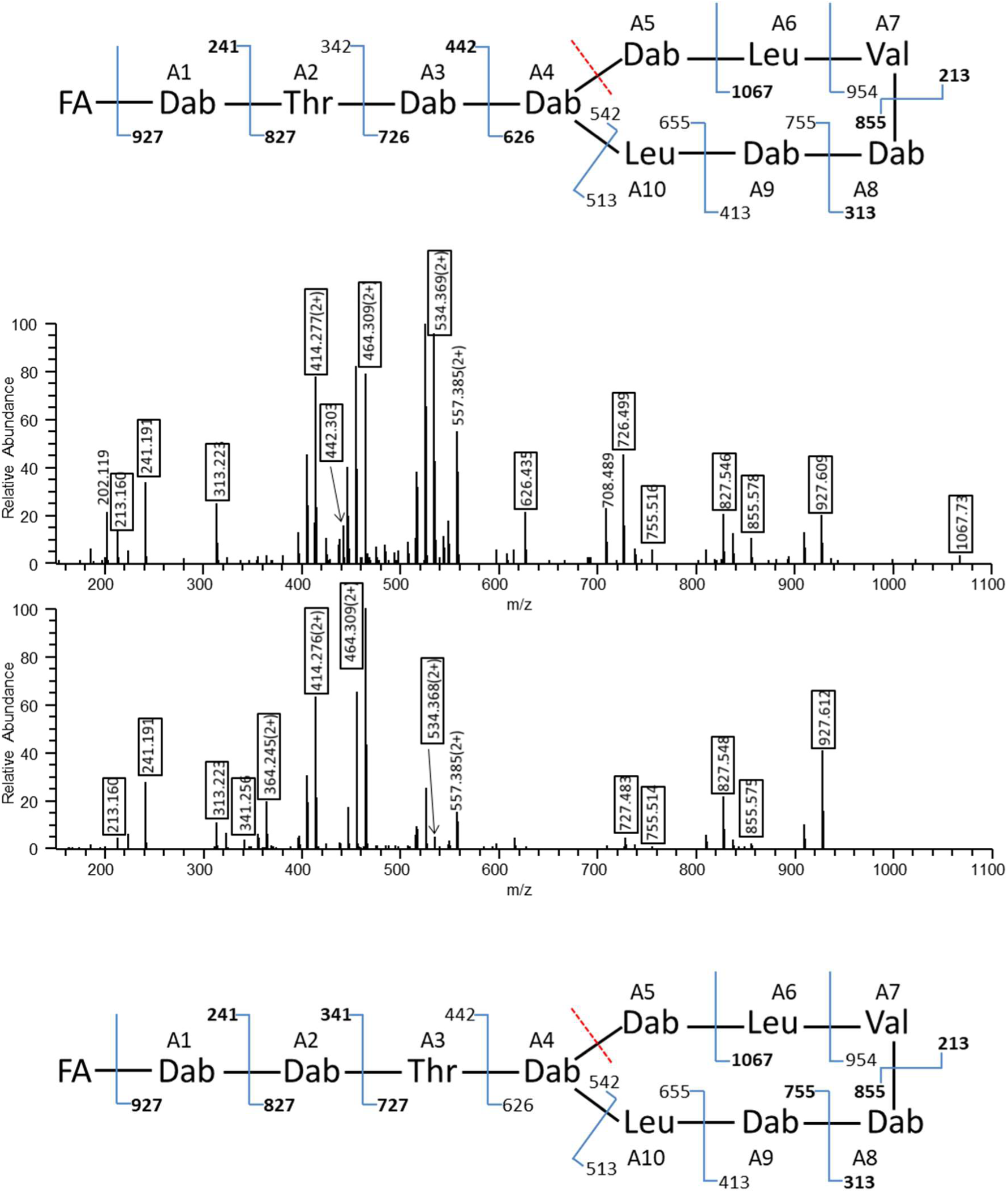
CID spectra of synthetic polymyxins (m/z 584.402 (2+)). Both peptides are N-capped with a C9-fatty acyl group (FA) and feature two Leu’s and one Val in the cyclic region. Top panel (C91): canonical N-terminal sequence stretch of Dab-Thr-Dab; Bottom panel (C93): N-terminal sequence is Dab-Dab-Thr. Potential peptide bond fragment ion series starting from the N-terminus or A5|A6 cleavage site are listed along with the structures deciphered and the ones observed are highlighted in the spectra. Dab: 2,4-diaminobutyric acid.

To test the antimicrobial spectrum of the four synthetic polymyxins, standard bacterial growth tests were applied in microplates, as described in the Methods and Materials. Commercially available colistin (polymyxin E) was used as a standard of comparison, since it is available as a human therapeutic in most countries. MIC (minimum inhibitory concentration) values which indicate a change compared to colistin are shown in **Table 4**. (Strains upon which the synthetic polymyxins display no differential activity are shown in **Table S3**). The toxicity of the four synthetic polymyxins on mammalian cells was tested in comparison to that of colistin. Similarly to the colistin control, the tested compounds were practically not cytotoxic on normal liver and kidney cell cultures in pharmacologically relevant concentrations (**Table S4**).

**Table 4.**
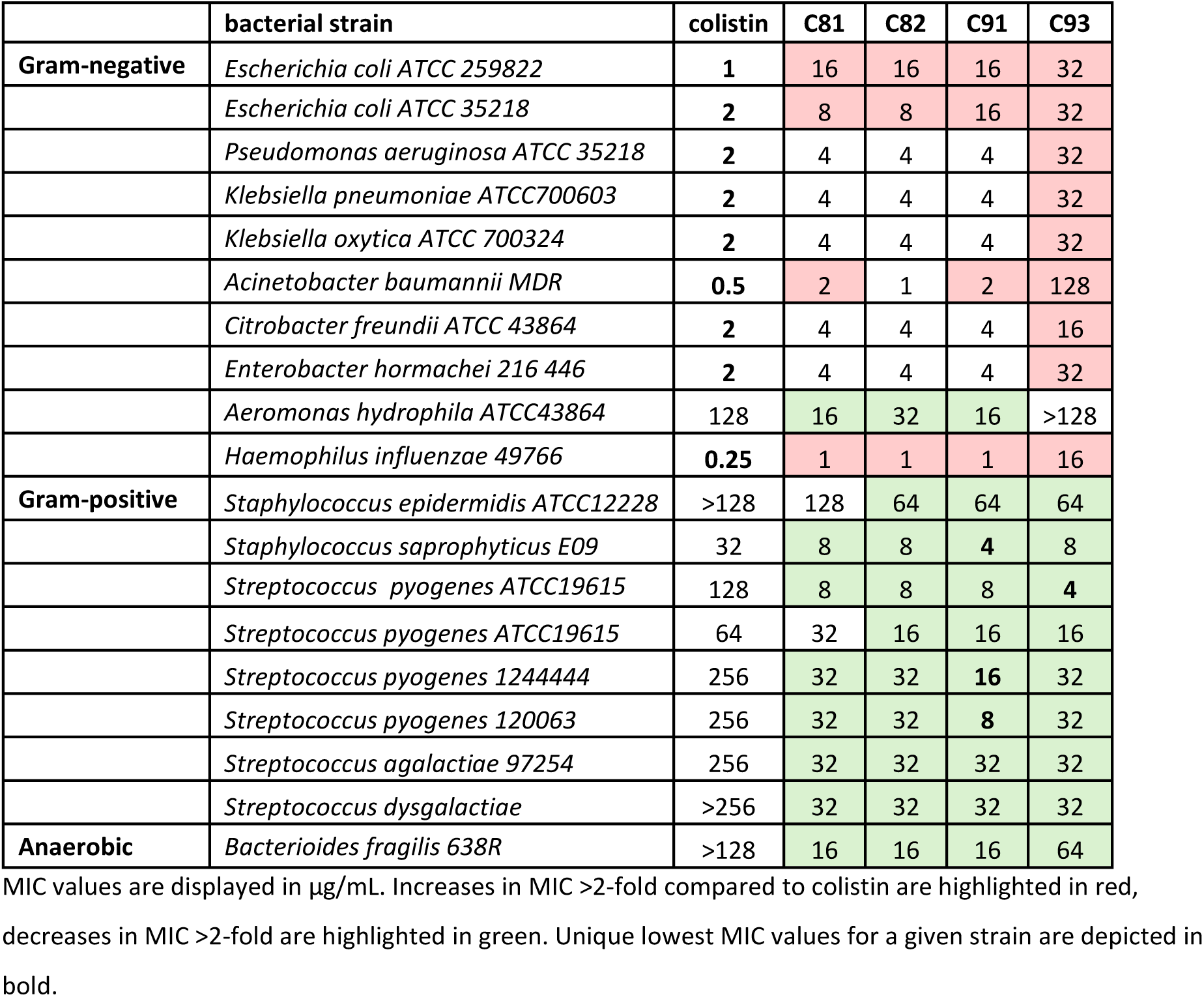
Minimal inhibitory concentrations (MICs) of colistin and the synthetic polymyxins, displayed against various human pathogens.

At this point we asked whether the N-terminal acyl chains are exclusively defined by the peptide sequence physically linked to them. To answer, we carefully re-inspected the MS data of the active fractions of ISO4. The ions representing both missing combinations of acyl chain + peptide (i.e. **C_9_-T1d6L3** and **C_8_-T1d6L2V1**, corresponding to MW of 1180.8 Da and 1152.8 Da, respectively) were detected within the active fractions of strain ISO4 (**Figure S7**), and MS/MS spectra provided evidence for the presence of these two peptides (**Figure S8**). This suggests that the N-terminal acyl chains are not defined by the peptide sequence, for all four acyl-peptide combinations were detectible.

Surprisingly, additional new polymyxin variants were also identified: the ion package representing the peptide MW 1152.8 proved to be a coeluting mixture of C8-T1d6L2V1 and C8-S1d6L3 (**Figure S9**, **top**), and the latter peptide was also found with the C9-acyl N-terminal capping (C9-S1d6L3, **Figure S9, bottom**). In the latter two, Ser-containing peptides, the Ser residue most likely resides in position A2 (**Figure S9**). These findings were somewhat unexpected considering that the prediction of the PARAS server indicated that the most probable substrate for A2 should be (E)-2-Butenyl-4-methyl-threonine (Bmt) (probability score 0.569) followed by Thr (0.431) while all other substrates, including Ser had a score of 0. We checked for the presence of Bmt using the calculated monoisotopic m/z values of C8-Bmt1d6L3 and C9-Bmt1d6L3 (m/z 412.624 and 417.296 representing the dominant charge state of z=3) and the Bmt reporter immonium ions (m/z 142.123 and 124.113) in the MS and MS/MS data of the ISO4 fractions, respectively; but no such components were detected (data not shown). The predictions of PARAS for A2 therefore proved to be only partially adequate, the ambiguity of the output nevertheless hinted at the promiscuity of this adenylation domain. The sequences of the putative natural polymyxins are summarized in **Table 5**.

**Table 5.** Putative sequences of natural polymyxins identified in this study. Variations are marked in red.

| Putative polymyxin sequence | Abbreviation* | Molecular weight (Da) |
| --- | --- | --- |
| octanoyl-Dab-Thr-dDab-cy(Dab-Dab-dLeu-Ile-Dab-Dab-Leu) | C8-T1d6L3 | 1166.8 |
| nonanoyl-Dab-Thr-dDab-cy(Dab-Dab-dLeu-Val-Dab-Dab-Leu) | C9-T1d6L2V1 | 1166.8 |
| octanoyl-Dab-Thr-dDab-cy(Dab-Dab-dLeu-Val-Dab-Dab-Leu) | C8-T1d6L2V1 | 1152.8 |
| nonanoyl-Dab-Thr-dDab-cy(Dab-Dab-dLeu-Ile-Dab-Dab-Leu) | C9-T1d6L3 | 1180.8 |
| octanoyl-Dab-Ser-dDab-cy(Dab-Dab-dLeu-Ile-Dab-Dab-Leu) | C8-S1d6L3 | 1152.8 |
| nonanoyl-Dab-Ser-dDab-cy(Dab-Dab-dLeu-Ile-Dab-Dab-Leu) | C9-S1d6L3 | 1166.8 |

\*The numbers after the letters denote the following: C- acyl chain length; T- number of Thr residues; d- number of Dab residues; L- number of Leu/Ile/Nle residues; V- number of Val residues; S- number of Ser residues

## Discussion

The constantly growing threat of antibiotic-resistant bacterial pathogens urges the discovery of novel antimicrobial therapeutics. This work describes an activity-based screening of culturable environmental bacteria in search of novel antimicrobial compounds. Besides the obvious aim of discovery, a further goal of this project was to obtain a snapshot on how much the repertoire of AMPs derived from such a screen are saturated by previously described compounds. Over 300 versatile environments were sampled, and culturing was supported by the use of multiple rich media both under aerobic and anaerobic conditions to improve the chances of success. Overall, nearly 10% of the samples turned out to contain a culturable microbe that could inhibit the growth of either of the two target bacteria. We continued with the activity-based HPLC separation of the extracts produced by axenic liquid cultures of the two most abundant genus within our study, the *Paenibacillus* and the *Brevibacillus* genus. The MS and MS/MS analyses revealed that in all active fractions, the antimicrobial effect was attributable to peptide-type molecules; polymyxins or closely related octapeptins from Paenibacilli and predominantly bogorols from Brevibacilli (data not shown). Of the 10 distinct polymyxins/octapeptins identified by their MS and CID spectra (**Table 3**), two promised to be novel polymyxin variants, this work was therefore dedicated to their analysis. Polymyxins, discovered in the 1940s, represent one of the earliest known classes of antimicrobial peptides (Peirano et al., 2011). Although their considerable side effects prevent their systemic application in everyday patient care today, they have found their place both as topical ointments and as last resort antibiotics to be used in cases of multi-resistant bacterial infections (Nation et al., 2015). Polymyxins are mostly effective against Gram-negative bacteria, having the ability to bind to the Lipid A component of their lipopolysaccharide (LPS), and ultimately leading to the permeabilization of both inner and outer membranes (Poirel et al., 2017). Polymyxins themselves are produced by Gram-positive bacteria, mostly belonging to the *Paenibacillus* genus (Storm et al., 1977). Structurally, they are usually decapeptides, comprising a C-terminal heptapeptide cycle preceded by an N-terminal linear tripeptide (Trimble et al., 2016). A nonpolar fatty acyl chain is usually connected to their N-terminus *via* an amide bond. The peptides are not directly encoded on the genome of the producer bacteria, but instead, are synthesized by huge multi-domain protein complexes, called non-ribosomal peptide synthases (NRPS). Their most commonly used variants are polymyxin B and polymyxin E (Colistin) (Evans et al., 1999), but hundreds of further sequence variants have also been described (J. Li et al., 2015). Due to the conserved nature of NRPS domains, it is possible to predict the peptide sequence of polymyxins from the DNA sequence of the NRPS gene cluster using antiSMASH, a well-established toolbox (Blin et al., 2023).

In our project, the output of the MS and CID analysis, the annotations of antiSMASH and the literature were all used to define the structure of the polymyxins detectable in the extract of *Paenibacillus* strain ISO4. The antiSMASH server identified only a single NRPS capable of polymyxin synthesis in the genome, and notoriously, the specificity of the adenylation domain of module 7 was either undefined or was ambiguous during repeated analyses. The cessation of the production of both novel polymyxins in our *pmxE* mutant *Paenibacillus* strain nevertheless mapped the production of both AMPs to this single NRPS-type gene cluster. This suggests that enzyme promiscuity is responsible for the sequence variation seen in the novel polymyxins, standing in agreement with earlier reports describing the promiscuous substrate specificities of the adenylation domains of modules 3, 6 and 7 (Galea et al., 2017).

Our current claim is therefore that the two polymyxins represented by the dominant ion of m/z 389.936(3+) are most likely **octanoyl-Dab-Thr-dDab-cy(Dab-Dab-dLeu-Ile-Dab-Dab-Leu),** and **nonanoyl-Dab-Thr-dDab-cy(Dab-Dab-dLeu-Val-Dab-Dab-Leu).** These correspond to **C_8_-T1d6L3** and **C_9_-T1d6L2V1** in **Table 3**, and both have the molecular weight (MW) of 1166.8 Da. To the best of our knowledge, this is the first time the natural production of full length polymyxins harboring three aliphatic residues at positions 6, 7 and 10 were described. Besides these two dominant polymyxins, four minor AMPs were also detected. These are most likely to be **nonanoyl-Dab-Thr-dDab-cy(Dab-Dab-dLeu-Ile-Dab-Dab-Leu), octanoyl-Dab-Thr-dDab-cy(Dab-Dab-dLeu-Val-Dab-Dab-Leu), octanoyl-Dab-Ser-dDab-cy(Dab-Dab-dLeu-Ile-Dab-Dab-Leu), and nonanoyl-Dab-Ser-dDab-cy(Dab-Dab-dLeu-Ile-Dab-Dab-Leu).** Again, defining their exact structure was beyond the scope of this study due to the limitations of mass spectrometry and the lack of their sufficient purity and yield required for NMR studies. Synthetic polymyxins harboring Ser at A2 have been described multiple times (Kline et al., 2001; J. Li et al., 2021; Vaara & Vaara, 2010), however we have not seen it reported in natural samples before. Interestingly, polymyxins with Ser at A3 were claimed to be produced by a strain of *Paenibacillus polymyxa* (Galea et al., 2017).

In this work, the information collected on natural polymyxins was used to design novel synthetic variants. The four sequences (**Figure 6**) were chosen to include Leu, Ile and Val at position A7, test a C8 and a C9 linear acyl chain on the N-terminus, and investigate the consequences of swapping A2 and A3 on the antimicrobial effect and the CID spectrum. (Originally, a **C_9_-T1d6L2V1 polymyxin with** the Val at position A6 was also ordered, but the shipped AMP had lacked cyclization, as revealed by the MS analysis. It displayed a marginal antimicrobial effect and was not included in the study.) Overall, 29 human pathogens representing 21 species were challenged with the four synthetic polymyxins and colistin (**Table 4, S3**). A decreased MIC (i.e. increased potency) of either synthetic polymyxin compared to that of colistin could be detected against six species. Of these, *Aeromonas hydrophila* is an opportunistic pathogen in humans and poses a significant threat to global food safety due to its ability to cause devastating diseases in farmed and wild fish (Jeamsripong et al., 2025). *Bacterioides fragilis* is an important anaerobic member of the normal human gut flora, in case of bowel perforation however, it can cause serious intra-abdominal infections, abscesses, and sepsis (Wexler, 2014). *Staphylococcus epidermidis* constitutes a significant component of the normal microflora in humans, but is a potential cause of nosocomial infections (Götz et al., 2006).

*Staphylococcus saprophyticus* is the second most common cause of uncomplicated urinary tract infections, after *Escherichia coli* (Flores-Mireles et al., 2015). *Streptococcus dysgalactiae* is responsible for pharyngitis connected with consumption of contaminants in dairy products (Hardie & Whiley, 2006). And finally, *Streptococcus pyogenes* is a frequently detected, versatile pathogenic bacterium responsible for numerous diseases ranging from pharyngitis to rheumatic fever and glomerulonephritis (Hardie & Whiley, 2006). The tested strains of *S. saprophyticus* and *S. pyogenes* were inhibited by our novel synthetic polymyxins in clinically relevant concentrations.

The most valuable synthetic polymyxin of our tests was C82, surpassing colistin against ten pathogens, and underperforming against three. Approaching from a different aspect, the C91 compound turned out to display the unique lowest MIC against the highest number of pathogens (three) tested. The MIC test also clarified that changing the N-terminal Ac-Dab-Thr-Dab sequence to Ac-Dab-Dab-Thr (as in C93) increases MIC (reduces potency) in nine cases, all of which are Gram-negative pathogens. In case of targeting Gram-positives however, C93 displays the same degree of improvement over colistin as do the other three synthetic polymyxins. The Ac-Dab-Dab-Thr sequence pattern is therefore not recommended in generalist polymyxin design but could be a rational choice if the spectrum is to be narrowed to Gram-positives. The differences in MIC values between C81, C82 and C91 were maximum 2-fold for any targeted pathogen. This is not considered significant by convention; we can therefore treat them as a highly similar group of polymyxins. This stands in line with earlier structure-activity relation studies: exchanging Leu-7 for Ile or Val was found to have no substantial effect on the antimicrobial spectrum of polymyxin E (Cui et al., 2018; Kline et al., 2001).

Akin to our result in C93, the substitution of Thr-2 with Dab in a polymyxin B significantly increased MIC to 128 µg/mL against *P. aeruginosa* PA01 (J. Li et al., 2021). This is also similar to the reports of Cui et al. (Cui et al., 2020) in which a similar substitution increased the MIC against Gram-negatives. An important exception to this trend of decreased potency against Gram-negatives was the case when targeting *Moraxella catarrhalis* ATCC 25238, where C93 displayed activity similar to colistin (4 µg/mL)(**Table S3**). The unique activity of C93 against this Gram-negative pathogen, despite the Thr-to-Dab substitution at A2 suggests that the activity observed for polymyxin analogues may depend on the nature of individual target bacteria’s LPS composition. For example, the LPS of *M. catarrhalis* has been reported to lack the O-polysaccharide chain (Holme et al., 1999).

Overall, our project has so far yielded four novel synthetic polymyxin sequences, which were inspired by six newly discovered natural polymyxin variants. The final evaluation of these results boils down to the question whether initiating a classical functional screen of environmental samples is a worthwhile endeavor to discover novel antimicrobial compounds. Based on the numbers, one might deduce that screening over 300 environmental samples for the discovery of novel variants of a known AMP is an unprofitable investment. We argue however the opposite, since two new subclasses of natural AMPs could be discovered this way: i) full-length polymyxins that contain three aliphatic residues within their ring, at positions A6, A7 and A10, and ii) full-length polymyxins that contain a Ser residue at position A2. The four polymyxins designed based on the structures hinted at by the MS/MS data, antiSMASH and literature are far from exhausting the possibilities of the true structures potentially present within the single producer strain this worked focused on. We therefore agree with earlier declarations that naturally derived samples, and more specifically, soil bacteria still provide a vast untapped source of biomolecules potentially useful for healthcare applications.

## Acknowledgements

The authors thank Gábor Draskovits and Dr. Jangir Pramod Kumar for donating certain soil samples, Dr. Josef Altenbuchner for providing the pJOE8999 plasmid, and Prof. Katalin F. Medzihradszky for consulting. We sincerely thank scientific and non-scientific staff members of the Institute of Biochemistry, HUN-REN Biological Research Centre, Szeged, Hungary, and the staff of the sequencing, proteomics, and metabolomics core facility of the institute.

