## Supplementary material for "Novel Synthetic Polymyxin Variants Inspired by Newly Uncovered Natural Sequences Explored as Potential Antibiotics"

### Supplementary tables

**Table S1.** Oligonucleotides used in this study

| primer name | sequence | function |
|---|---|---|
| Xba26_16Up_F | GTCTAGACGCAAGCCCAACG | primers for contructing pJOE26_16 |
| 26_16Up_R | CCGAAACTCTGTACTTCGTCAGCCAG | primers for contructing pJOE26_16 |
| 26_16Dn_F | CGAAGTACAGAGTTTCGGGCGAG | primers for contructing pJOE26_16 |
| Xba26_16Dn_R | GTCTAGATCTCCGGTCGGTTC | primers for contructing pJOE26_16 |
| sg26_16F | TACGGACAGCACCGACGTTCCGGT | primers for contructing pJOE26_16 |
| sg26_16R | AAACACCGGAACGTCGGTGCTGTC | primers for contructing pJOE26_16 |
| 27F | AGAGTTTGATCCTGGCTCAG | primers for amplifying bacterial rDNA |
| 1492R | GGTTACCTTGTTACGACTT | primers for amplifying bacterial rDNA |
| pJOEhomF | gatgccactcttatccatcaatcc | primers for detecting pJOE26_16 |
| pJOEhomR | tttgttcagaacgctcggttg | primers for detecting pJOE26_16 |
| 26_16checkA | AACGGGAGTGTGTCGATGTG | primers for detecting the integration of pJOE26_16 |
| 26_16checkB | CGCAAAGGCCAAGACATCTC | primers for detecting the integration of pJOE26_16 |
| 26_16checkC | ACTTCTCGGCAGTCAGATCC | primers for detecting the integration of pJOE26_16 |
| 26_16checkD | TTCCAGCATGGTACGTGGAG | primers for detecting the integration of pJOE26_16 |

**Table S2.** Composition of the Micronutrient Booster solution

| component | concentration (mg/mL) |
|---|---:|
| $FeSO_4 \cdot 7H_2O$ | 2 |
| $ZnSO_4 \cdot 7H_2O$ | 2.23 |
| $MgSO_4 \cdot 7H_2O$ | 0.4 |
| $MnSO_4 \cdot H_2O$ | 0.31 |
| $CuSO_4 \cdot 5H_2O$ | 0.25 |
| $(NH_4)_6Mo_7O_{24} \cdot 4H_2O$ | 0.19 |
| $H_4NO_3V$ | 0.12 |
| $H_3BO_3$ | 0.1 |
| NaF | 0.09 |
| $CoCl_2 \cdot 6H_2O$ | 0.025 |
| $CrCl_3 \cdot 6H_2O$ | 0.018 |
| $Na_2SeO_3$ | 0.012 |

**Table S3**. Minimal inhibitory concentrations (MICs) of colistin and the synthetic polymyxins, displayed against various human pathogens

| | **bacterium** | **colistin** | **C81** | **C82** | **C91** | **C93** |
|---|---|---|---|---|---|---|
| **Gram-negative** | *Burkholderia cepacea ATCC 25416* | >128 | >128 | >128 | >128 | >128 |
| | *Moraxella catarrhalis ATCC25238* | 4 | **2** | **2** | **2** | 4 |
| **Gram-positive** | *Staphylococcus aureus ATCC25923* | >128 | >128 | >128 | >128 | >128 |
| | *Staphylococcus aureus ATCC29213* | >128 | >128 | >128 | >128 | >128 |
| | *Staphylococcus aureus ATCC12493 (MRSA)* | >128 | >128 | >128 | >128 | >128 |
| | *Staphylococcus saprophyticus ATCC BAA-750* | 8 | 8 | **4** | 8 | **4** |
| | *Enterococcus faecalis ATCC29212* | >128 | >128 | >128 | >128 | >128 |
| | *Enterococcus faecium ATCC700721* | >128 | >128 | >128 | >128 | >128 |
| | *Streptococcus agalactiae 95470* | >256 | 256 | 256 | 256 | >256 |
| | *Streptococcus pneumoniae ATCC49619* | >256 | >256 | >256 | >256 | >256 |

MIC values are displayed in µg/mL.

**Table S4.** The $IC_{50}$ values of various polymyxin peptides measured on three mammalian cell lines

| | HepG2 | Hek293 | HRPTEpC |
|---|---|---|---|
| C81 | >100 µg/ml | >100 µg/ml | 50 µg/ml |
| C82 | >100 µg/ml | >100 µg/ml | 70 µg/ml |
| C91 | >100 µg/ml | >100 µg/ml | 50 µg/ml |
| C93 | >100 µg/ml | >100 µg/ml | 40 µg/ml |
| Colistin | >100 µg/ml | >100 µg/ml | 90 µg/ml |

**Supplementary figures**

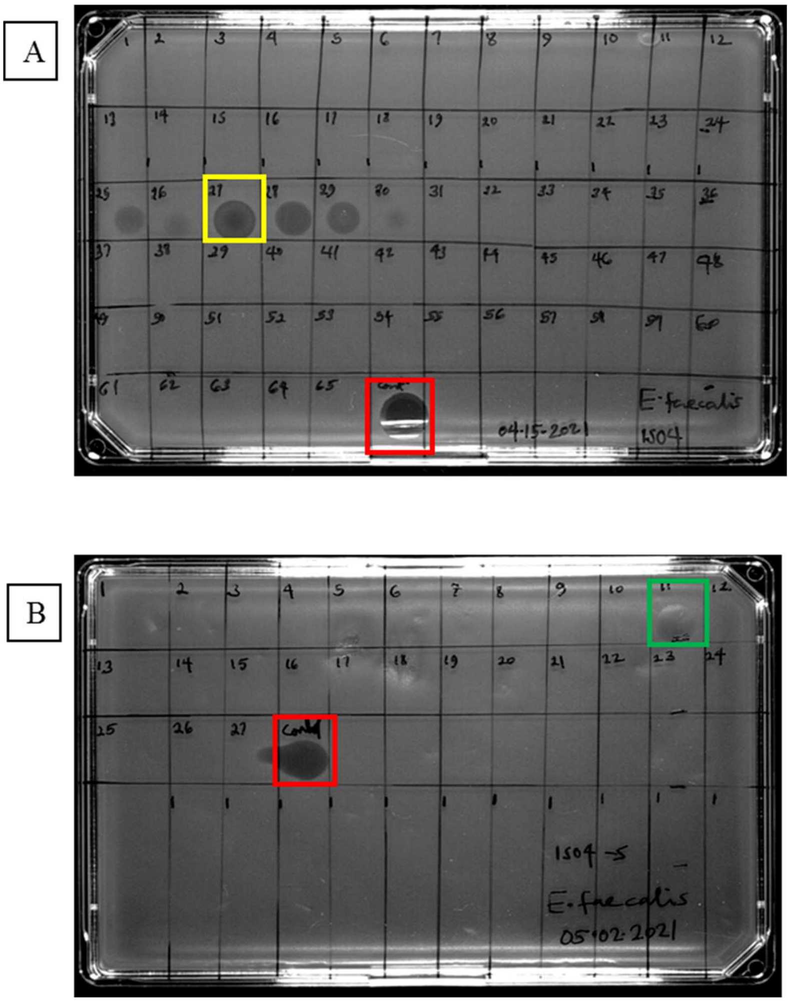


**Figure S1.** Antibacterial activity of HPLC fractions from *P. barciconensis* (ISO4) targeting *E. faecalis* (A): after crude extract separation (B): after second dimension separation. Fraction number 27 in the yellow square coded (ISO4_27) indicates the active fraction chosen for the second round of HPLC separation. Fraction number 11 in the green square was submitted for MS analysis. Fractions in the red square indicate the positive control, i.e. the injected sample of the respective separation plated directly

**Identified secondary metabolite regions using strictness 'relaxed'**

| Region | Type | From | To | Most similar known cluster | | Similarity |
|---|---|---|---|---|---|---|
| Region 1.1 | lassopeptide | 220,767 | 244,736 | paeninodin | RiPP | 60% |
| Region 6.1 | lassopeptide | 17,893 | 40,007 | | | |
| Region 6.2 | ranthipeptide | 66,439 | 88,065 | | | |
| Region 6.3 | lassopeptide | 100,210 | 115,966 | | | |
| Region 7.1 | T3PKS | 1 | 30,588 | | | |
| Region 19.1 | transAT-PKS, T1PKS, NRPS | 18,540 | 81,784 | xenocoumacin 1 / xenocoumacin II | NRP + Polyketide:Modular type I | 21% |
| Region 21.1 | NRPS-like | 1 | 25,781 | | | |
| Region 23.1 | lanthipeptide-class-iv | 19,172 | 41,922 | | | |
| Region 26.1 | NRPS | 1 | 71,844 | colistin A / colistin B | NRP | 100% |
| Region 29.1 | proteusin | 20,842 | 41,090 | | | |
| Region 44.1 | siderophore | 31,258 | 49,240 | | | |
| Region 76.1 | lanthipeptide-class-i | 1 | 15,242 | | | |
| Region 94.1 | terpene | 1 | 15,737 | carotenoid | Terpene | 33% |

**Figure S2.** Identification of biosynthetic gene clusters in *Paenibacillus* strain ISO4 with antiSMASH

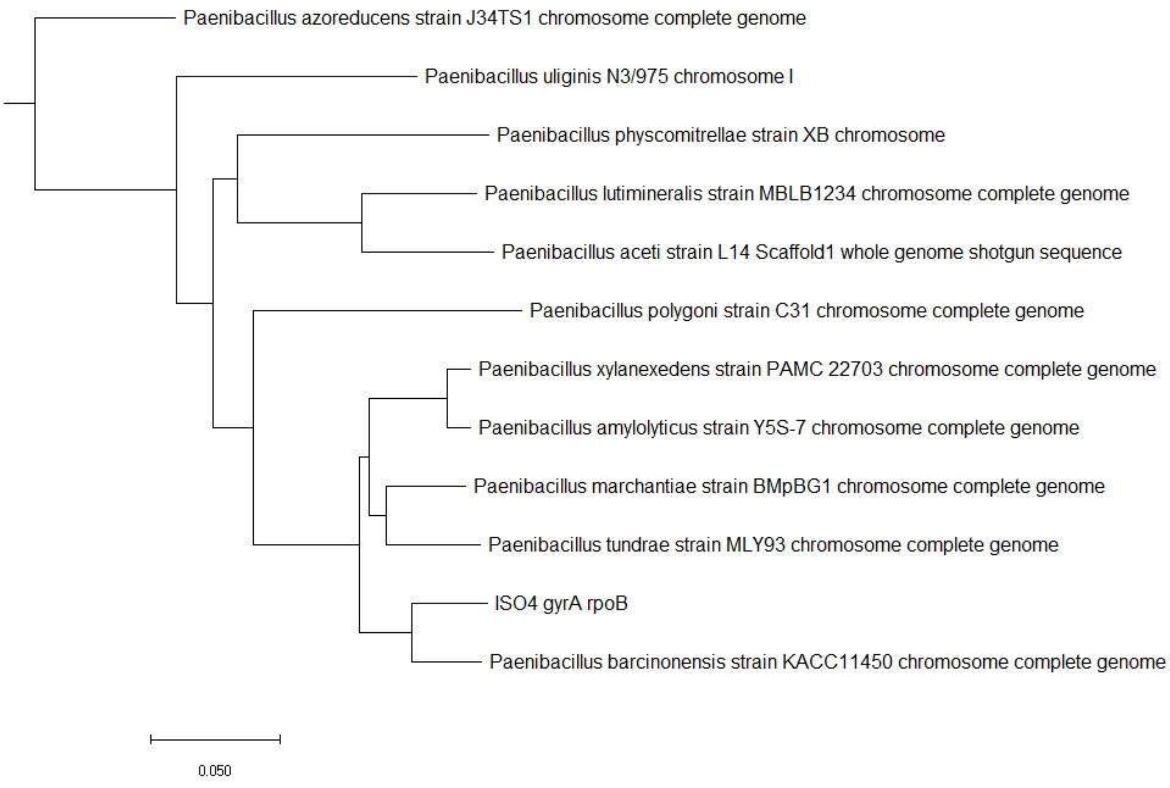


**Figure S3**: Neighbor-joining phylogenetic tree of *P. barciconensis* (ISO4) constructed based on concatenation of *gyrA* and *rpoB* gene sequences.

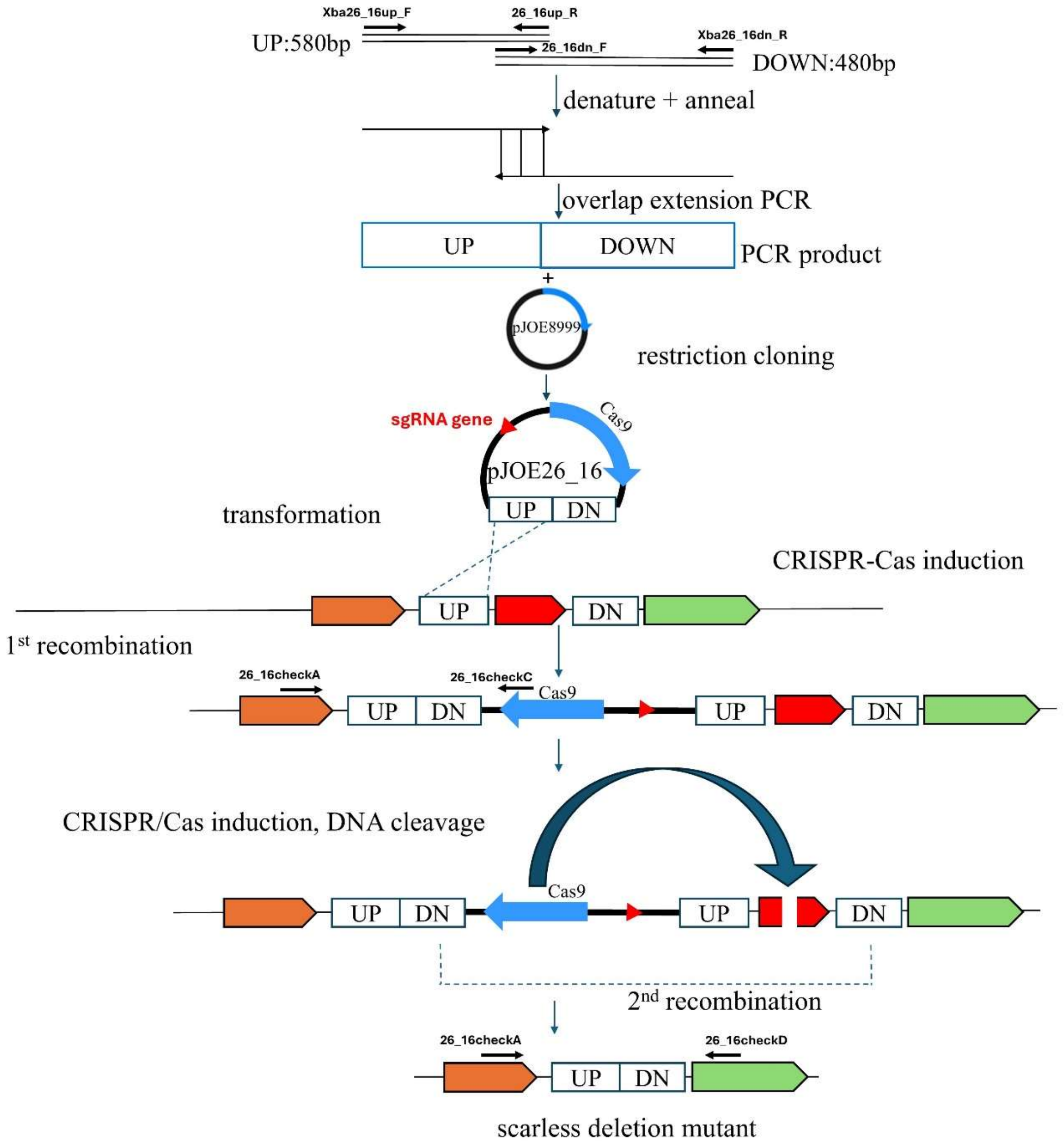


**Figure S4**. Schematic representation of the method used to delete a 1.8 kbp segment of gene ctg26_16 from the genome of *Paenibacillus* strain ISO4

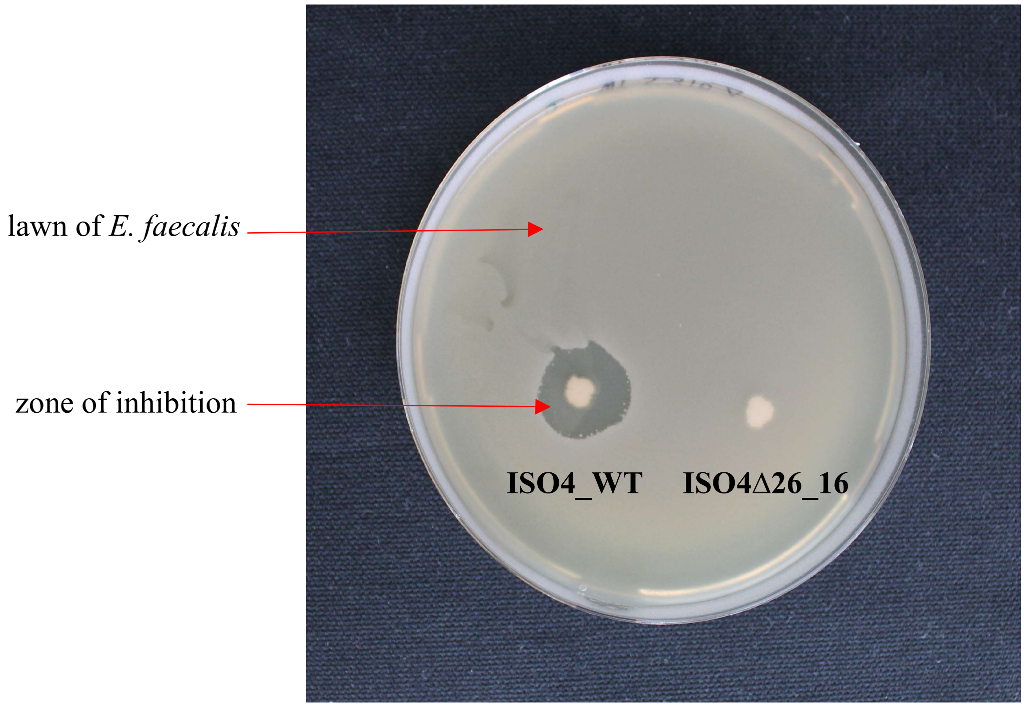


**Figure S5.** Comparison of antibacterial activity of wild–type *P. barciconensis* (ISO4) and deletion mutant (ISO4Δ26_16) by soft-agar overlay assay targeting *E. faecali*s.

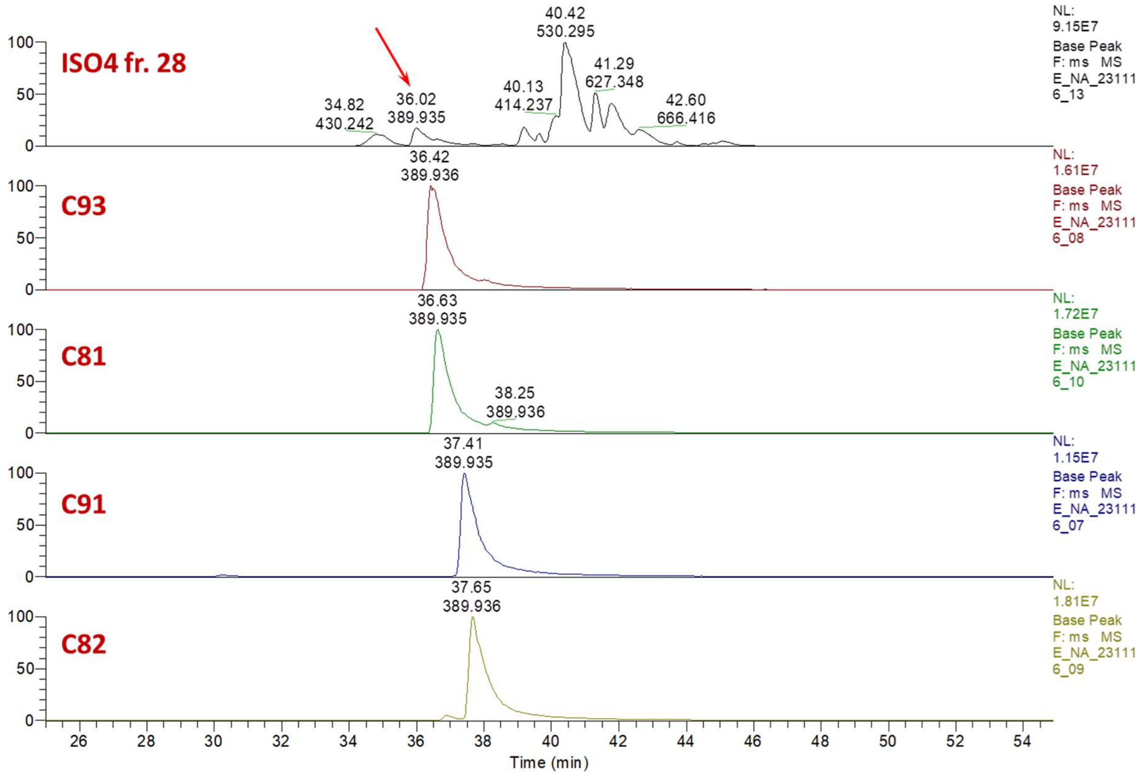


**Figure S6. LC-MS/MS BPI profiles of the active fraction of ISO4 and the synthetic polymyxins.** The natural polymyxin peptide in ISO4 is indicated by an arrow. For structures of the synthetic peptides, see Figure 7. All traces were smoothed using a 3-point boxcar smoothing algorithm.

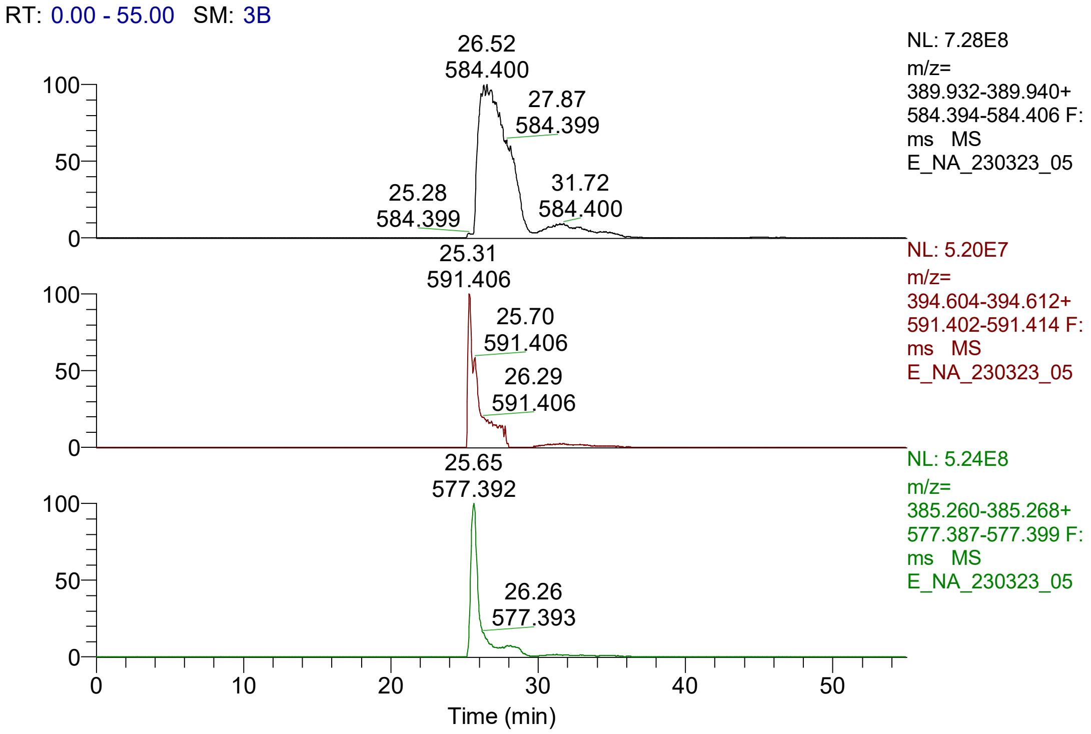


**Figure S7. Extracted ion chromatograms (XIC) of different polymyxin peptides, generated from the LC-MS/MS data of the active fraction of isolate ISO4.**

Top panel: XIC for polymyxin compositions C8-T1d6L3 or C9-T1d6L2V1 (m/z 389.936 and 584.764, representing the triply and doubly charged peptides, respectively)
Middle panel: XIC for polymyxin composition C9-T1d6L3 (m/z 394.608 and 591.408, representing the triply and doubly charged peptide, respectively)
Bottom panel: XIC for polymyxin composition C8-T1d6L2V1 (m/z 385.264 and 577.393, representing the triply and doubly charged peptide, respectively)
All XIC traces were generated with 10 ppm mass error. C8: octanoyl, C9: nonanoyl, d: 2,4-diaminobutiric acid, T: threonin, L: leucin, V: valine

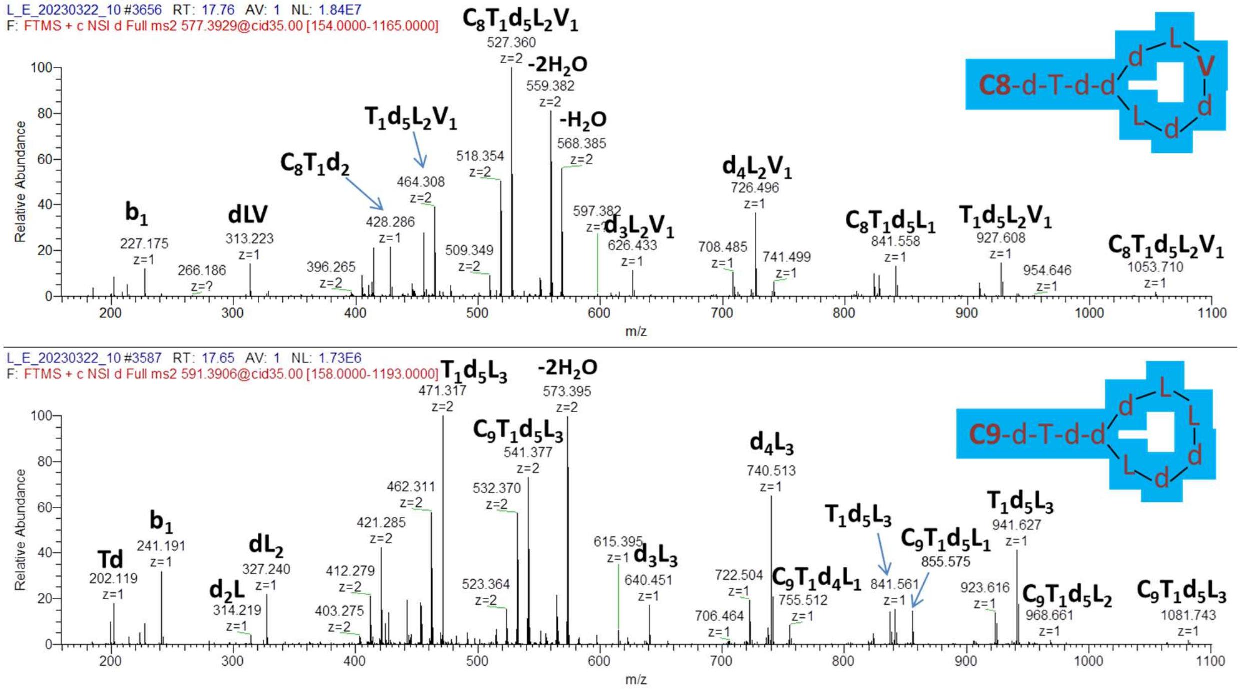


**Figure S8. MS/MS spectra of precursor ions corresponding to polymyxin compositions of C8-T1d6L2V1 (top; *m/z* 577.39, 2+) and C9-T1d6L3 (bottom; *m/z* 591.39, 2+).**

C8: octanoyl, C9: nonanoyl, d: 2,4-diaminobutiric acid, T: threonin, L: leucin, V: valine, -$H_2O$: water loss of the precursor ion
MS/MS data were acquired in the linear ion trap of on Orbitrap Lumos Fusion Tribrid mass spectrometer.

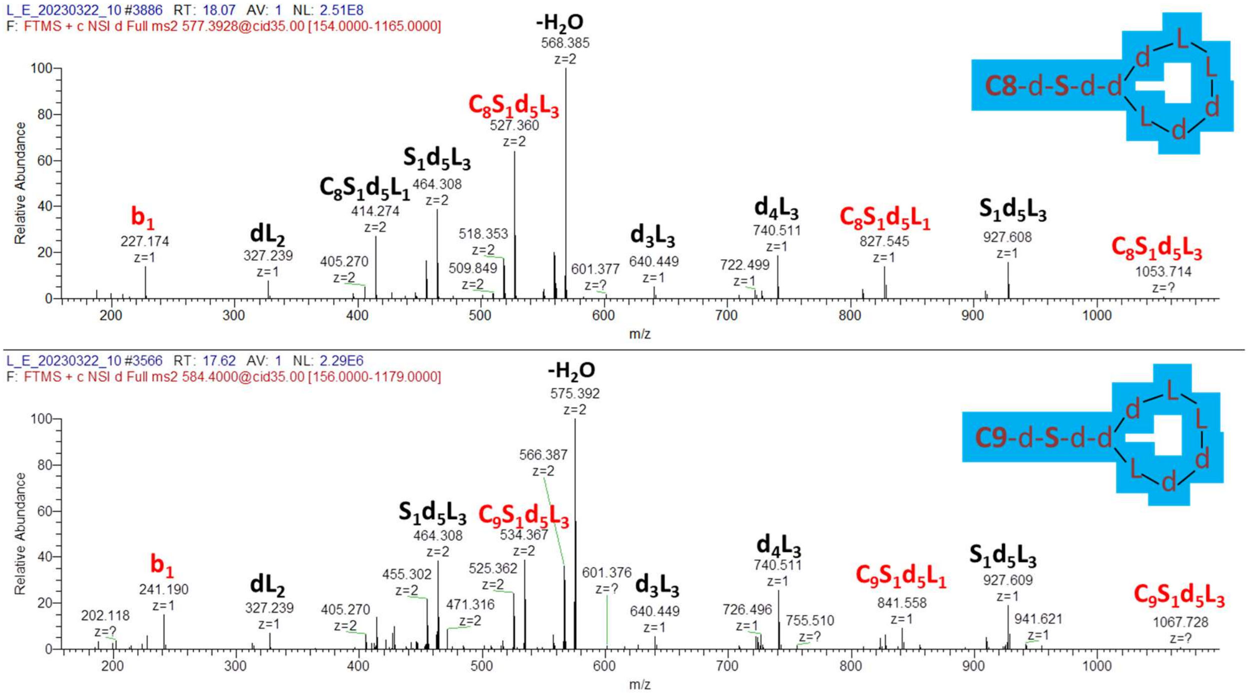


**Figure S9. MS/MS spectra of precursor ions corresponding to polymyxin compositions of C8-S1d6L3 (top; *m/z* 577.39, 2+) and C9-S1d6L3 (bottom; *m/z* 584.40, 2+).**

These peptides differ only in the N-terminal acyl group as indicated by the abundant $b_1$ ions at m/z 227 and 241 and additional fragment ions with intact peptide N-terminus (highlighted in red). Both peptide harbor the same ring structure consisting of 4 Dab and 3 Leu residues as evidenced by the fragment ion of m/z 740.51 present in both spectra. The presence of Ser instead of conventional Thr as A2 is concluded based on the m/z of the precursor ions and supporting fragment ions such as m/z 927.61 present in both spectra.
C8: octanoyl, C9: nonanoyl, d: 2,4-diaminobutiric acid, S: serine, L: leucin, -$H_2O$: water loss of the precursor ion
MS/MS data were acquired in the linear ion trap of on Orbitrap Lumos Fusion Tribrid mass spectrometer.

**Supplementary discussion**

We observed some discrepancies in the fragmentation pattern of the extracted polymyxin peptides. CID analysis performed in ion traps usually leads to single bond cleavages, as the fragment ions' m/z values are different from the precursor ion, and thus, they are not activated therefore do not fragment further. As a result, internal fragment ions are usually not generated. Cyclic peptides represent a special case, since a bond cleavage within the ring will yield a mass identical to that of the precursor ion, and thus, further fragmentation may occur. However, in the earlier and later eluting polymyxin variants identified in this study (**Figure 1**), an abundant ion pair, m/z 827 and 841 cannot be assigned following the above rule, as these ions could be annotated only when the cleavage of 3 peptide bonds (A1|A2, A4|A5 and A5|A6) is considered. Similar atypical fragmentation in polymyxin ion trap CID spectra has been presented but not addressed in earlier publications (Govaerts, Orwa, Schepdael, et al. 2002; Govaerts, Orwa, Van Schepdael, et al. 2002). One possible explanation for the observed fragmentation is that the A2 and A3 amino acids are swapped (from Thr-Dab to Dab-Thr). In order to test this, we ordered the synthesis of polymyxins C91 and C93, carrying FA-Dab-dThr-dDab or FA-Dab-dDab-dThr on their N-termini, respectively, and compared their fragmentation patterns. (For simplicity of stereoisomer positioning, we chose both A2 and A3 to be D-amino acids in both variants.) We observed the same atypical fragmentation for these synthetic peptides as well: e.g. fragment ion at m/z 727 for the FA-Dab-Dab-Thr variant can only be explained by 3 peptide bond cleavages (A1|A2, A4|A5 and A5|A6) (**Figure 7**). We therefore concluded that the fragment ions observed in the natural sample cannot be taken as evidence for the non-canonical N-terminal pattern (Ac-Dab-Dab-Thr). Presently we cannot offer any explanation for the observed fragmentation pattern detailed above. Considering that the Dab-Dab-Thr version shows inferior antibiotic effect (**Table 4**), we did not pursue this issue further.

Govaerts, Cindy, Jennifer Orwa, Ann Van Schepdael, Eugène Roets, and Jos Hoogmartens. 2002. “Characterization of Polypeptide Antibiotics of the Polymyxin Series by Liquid Chromatography Electrospray Ionization Ion Trap Tandem Mass Spectrometry.” *Journal of Peptide Science* 8(2): 45–55. doi:10.1002/psc.367.

Govaerts, Cindy, Jennifer Orwa, Ann Van Schepdael, Eugène Roets, and Jos Hoogmartens. 2002. “Liquid Chromatography-Ion Trap Tandem Mass Spectrometry for the Characterization of Polypeptide Antibiotics of the Colistin Series in Commercial Samples.” *Journal of chromatography. A* 976(1–2): 65–78. doi:10.1016/s0021-9673(02)00375-8.
